# Electron counting enables cryo-electron ptychography for near-atomic-resolution cryo-electron microscopy

**DOI:** 10.64898/2026.07.25.736262

**Authors:** Shouqing Li, Bo Shen, Zhaoyi Yan, Junjiang Liu, Chenxi Tang, Zhao Wang, Zhi Deng, Xueming Li

**Affiliations:** Beijing Frontier Research Center for Biological Structure, Beijing 100084, China; Shenzhen Medical Academy of Research and Translation, Shenzhen 518100, China; Key Laboratory for Protein Sciences of Ministry of Education, Tsinghua University, Beijing 100084, China; State Key Laboratory of Membrane Biology, Tsinghua University, Beijing 100084, China; Tsinghua-Peking Joint Center for Life Sciences, Beijing 100084, China; School of Life Sciences, Tsinghua University, Beijing 100084, China; Department of Automation, Tsinghua University, Beijing 100084, China; Key Laboratory of Particle & Radiation Imaging of Ministry of Education, Tsinghua University, Beijing 100084, China; Department of Engineering Physics, Tsinghua University, Beijing 100084, China

**Author notes:** **Correspondence should be addressed to:** Zhi Deng, Xueming Li. **These authors contributed equally to this work**: Shouqing Li, Bo Shen, Zhaoyi Yan, Junjiang Liu.

## Abstract

Cryo-electron ptychography is an emerging technique for studying radiation-sensitive biological specimens, developed from four-dimensional transmission electron microscopy (4D-STEM). Although ptychography has achieved ultrahigh resolution beyond conventional transmission electron microscopy limits for radiation-resistant samples, its application to frozen hydrated biological specimens currently remains at sub-nanometer resolution. Here we overcome this limitation by implementing electron counting with a hybrid-pixel detector, establishing key technical foundations for near-atomic-resolution cryo-ptychography. This counting approach significantly improves weak signal detection in convergent-beam electron diffraction, enabling ptychographic reconstruction at doses below 1 e^−^/Å². Additionally, we found beam-induced motion is effectively eliminated within single scans, suggesting conventional cryoEM’s dose-fractionation approach may need reevaluation. Demonstrating high contrast under both low-dose and tilted conditions, along with achieving 3.59 Å resolution for the ∼700 kDa T20S proteasome, we validate cryo-electron ptychography as a viable general imaging modality that could complement or surpass conventional phase-contrast cryoEM methods.

## Introduction

Scanning transmission electron microscopy (STEM) and phase-contrast imaging are two core imaging methods in transmission electron microscopy. In materials science, STEM and four-dimensional STEM (4D-STEM) (*1*) have developed into mainstream imaging tools, among which ptychography has received widespread attention in recent years as a computational imaging approach that surpasses the physical resolution limit of transmission electron microscopes (*2–4*) and supports lensless imaging. By contrast, phase-contrast imaging still dominates in cryo-electron microscopy (cryoEM). With advancements in hybrid-pixel detector technology, low-dose ptychography has become achievable. Its integration into cryo-electron microscopy, termed cryo-electron ptychography, holds significant promise and may present opportunities to address critical bottlenecks in cryoEM. However, the application of ptychography to biological specimens remains constrained to sub-nanometer resolution levels for single-particle analysis (SPA) (*5*, *6*), thus currently unsuitable for routine cryoEM implementation.

Ptychography has demonstrated superior dose efficiency by directly utilizing the full diffraction intensity for phase retrieval, thereby fully exploiting the scattering information from every incident electron. This dose efficiency manifests as enhanced contrast and resolution. Both numerical simulations (reaching approximately 5 e^−^/Å²) (*7*) and experimental studies on frozen hydrated virus specimens (*8*) confirm that ptychography delivers superior image contrast and signal-to-noise ratio (SNR) compared to conventional cryoEM phase contrast imaging, even when the latter employs higher electron doses. For thick specimens, this advantage is further amplified. Tilt-corrected bright-field (tcBF) imaging, which employs the same data acquisition method as ptychography, has demonstrated significant contrast enhancement for half-micrometer-thick cellular specimens, highlighting the distinct advantage of convergent-beam diffraction (*9*). In terms of resolution, current studies on SPA using cryo-electron ptychography, referred to as ptychographic SPA, reveal that the convergence semi-angle influences both image contrast and resolution—smaller convergence semi-angles (e.g., 3 mrad) favor enhanced contrast, whereas larger convergence semi-angles (e.g., 7 mrad) enable higher resolution transmission. Nevertheless, the resolution currently reported for ptychographic SPA remains at the sub-nanometer level.

Direct electron detection and electron-counting technologies have enabled resolution breakthroughs in cryoEM through phase-contrast imaging under low-dose conditions (*10–12*). Optimizing detectors to maximize SNR should equally serve as a prerequisite for applying ptychography to biological specimens. Currently, most ptychography applications focus on high-dose scenarios in materials science, where the camera’s single-electron detection performance is often trivial. Extending ptychography to biological specimens faces at least two technical challenges: first, whether the detector can record convergent-beam electron diffraction (CBED) signals with high efficiency; and second, whether the phase retrieval can stably work on extremely sparse data. Existing hybrid-pixel detectors already have significantly higher single-electron sensitivity than the monolithic active pixel sensor (MAPS) detector commonly used in cryoEM, and can even implement electron counting directly at the detector. Furthermore, event-driven cameras can accurately record the amplitude and the arrival time of each electron, thereby resolving electron behavior more precisely. However, the practical performance of these detectors in cryo-electron ptychography still needs validation. Briefly, the key factors to achieve near-atomic-resolution remain unknown.

In this work, we implement electron counting based on a hybrid-pixel detector and apply it to cryo-electron ptychography, aiming to improve the recording performance of CBED patterns under extremely low dose conditions, thereby enhancing ptychography performance and further improving the resolution of ptychographic SPA. Using the T20S proteasome as a standard test sample, we systematically compare the performance differences of multiple electron-counting strategies, and investigate the effects of convergence semi-angle, irradiation dose and beam-induced motion. Meanwhile, we examine the low-dose imaging in tilted sample scenarios and discuss its feasibility for cryo-electron tomography (cryoET).

## Results

### Hybrid-pixel detector and electron-counting strategies

A high-performance hybrid-pixel detector is one of the key equipment for ptychography. For low-dose imaging of frozen hydrated biological specimens, high frame rate and accurate single-electron detection are critical metrics. High frame rate enables flexible control of the irradiation dose and implementation of electron counting to improve single-electron detection accuracy. In this work, a custom designed camera, named EMPIX was used (*13*). It consists of a 500 μm-thick silicon detector, with a pixel size of 150 μm × 150 μm and an array of 128 × 128 pixels. It supports a full-frame rate up to 50 kfps. The energy deposition behavior of 300 keV electrons was simulated in the detector, and accordingly different electron-counting strategies were designed and evaluated.

Electrons scatter randomly multiple times in the detector and deposit energy along their trajectories, causing charge sharing between adjacent pixels. This effect is the main factor affecting electron-counting accuracy and the design of subsequent counting strategies. The scattering and energy deposition of electrons in a 500-μm-thick silicon detector were simulated using the Monte Carlo method (*14*) and the statistics of the charge-sharing effect were quantitatively analyzed, as well as its impact on signal crosstalk between neighboring pixels. Among the 500 simulated events of 300 keV electrons (**Fig. 1A** and **B**), their trajectories can be distributed with a lateral range of 300 μm in radius, resulting in typical event clusters in 3-5 pixels (**fig. S1A**). The energy deposited in the pixel along the electron trajectory, reflected as the pixel readout amplitude, also varies considerably (**Fig. 1C** and **fig. S1B**). The amplitude value is dependent on the length of the trajectory covered by the pixel and the energy loss (dE/dx) of the electron. Intuitively, less energy is deposited in pixels at incident positions and with less trajectory coverage, while the highest energy is deposited at the end of trajectories due to the high dE/dx value. Before electron counting, we need to subtract background to identify pixel clusters affected by individual electron events, which causes pixels with small values to be neglected after background subtraction. Accordingly, the cluster sizes are shrunk to mostly 1-3 pixels in the simulation (blue and green bars in **Fig. 1D**), which is highly consistent with the statistics shown by experimental data (purple bars in **Fig. 1D**). The goal of electron counting is to find the initial incident positions of electrons to accurately restore the probability density of electron wave. However, such clustering distribution caused by the charge-sharing effect, combined with the low accuracy of the background threshold, ultimately introduces uncertainty in determining the incident position of electrons.

**Fig. 1.**
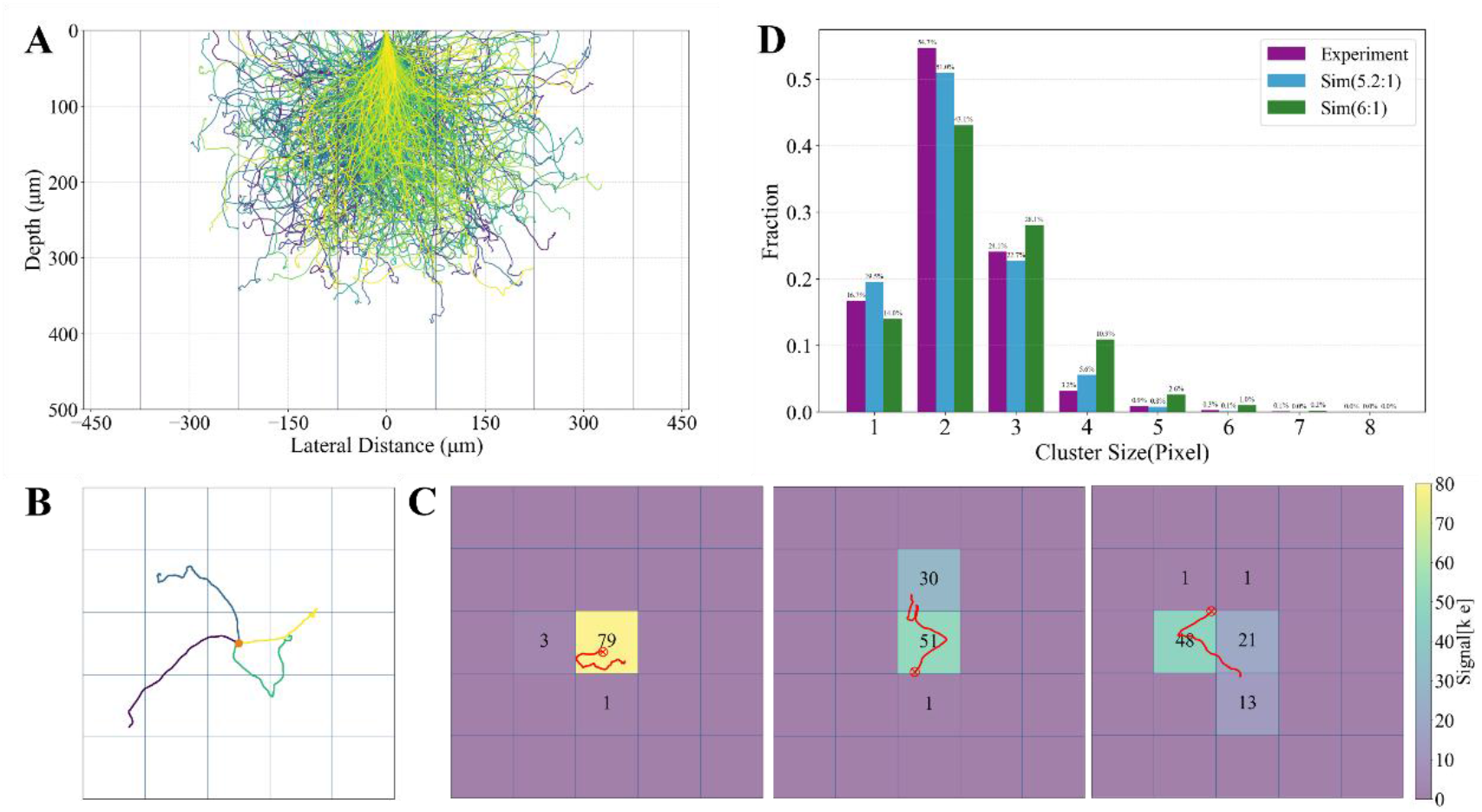
Electron detection simulation of EMPIX. (**A**) Simulated trajectories of 500 electrons with 300 keV energy in the silicon detector. Vertical solid lines demarcate pixel boundaries. (**B**) Projected trajectories of 4 simulated electron paths on the detector plane. (**C**) Simulated single-electron energy deposition across pixels. The red line denotes the electron trajectory, with the red marker ϒ indicating the incident position. The square lattice represents pixels, with color coding and numerical values inside indicating the corresponding pixel readout value. (**D**) Size distribution of pixel clusters affected by background subtraction. For simulated data, two statistical outcomes are presented with signal-to-threshold intensity ratios of 6:1 and 5.2:1. The former exhibits better agreement with experimental noise threshold, while the latter more closely approximates the experimental statistical distribution.

Based on the above understanding of the charge-sharing effect, we design several electron-counting strategies to address the uncertainty in determining the incident position of electrons. These strategies fall into two categories (see Methods). The first category selects only one pixel position in the cluster as the incident position, including strategies of selecting the strongest pixel (*S*^1*st*^), the third-strongest pixel (*S*^3*rd*^), and the weighted centroid pixel excluding the strongest pixel (*S^WCeM^*). The second category uses all pixels in the cluster, including strategies of setting all to 1 electron (*C*^1^) and setting all pixels to 1/*N* electron (*C*^1/*N*^), where *N* is the number of pixels in the cluster. We calculated the position residuals between these identified positions and the ground truth in the simulation (**fig. S1C**). Except for the strategy *S*^1*st*^ with larger deviation residuals, the other residuals are relatively close. The specific performance differences of the strategies require actual testing.

### Near-atomic-resolution single-particle reconstruction across counting strategies

Based on the counting strategies established above, we analyze the effects of different strategies on ptychographic SPA using the T20S proteasome. Experiments used a 5.10 mrad convergence semi-angle, with data acquired in a 328 nm × 328 nm field of view with a 350 × 350 scan grid. The number of readout electrons per scan position was approximately 1776, corresponding to a total detected electron dose of approximately 20.2 e^−^/Å² (see Methods). To implement electron counting, 20 consecutive CBED frames, termed the detector frames, were acquired at each scan position (**Fig. 2A**), with an average of 89 incident electrons identifiable per detector frame. The aforementioned electron-counting strategies were respectively applied to generate five datasets. Different strategies show slight differences in electron-counting positions (**Fig. 2B** and **fig. S2**). After counting, the 20 detector frames at each scan position were summed to produce the final CBED image (**Fig. 2C**), followed by ptychographic reconstruction of micrographs. Visual comparison shows that the micrographs with electron counting (**Fig. 2D**) are significantly better than those calculated directly from raw accumulation-mode data (**fig. S3A** and **B**). The accumulation-mode data acquired from a negative-stained T20S sample under high-dose conditions can be successfully reconstructed using ptychography (**fig. S3C**). These results highlight the crucial role of electron counting in enhancing camera performance at low-dose conditions.

**Fig. 2.**
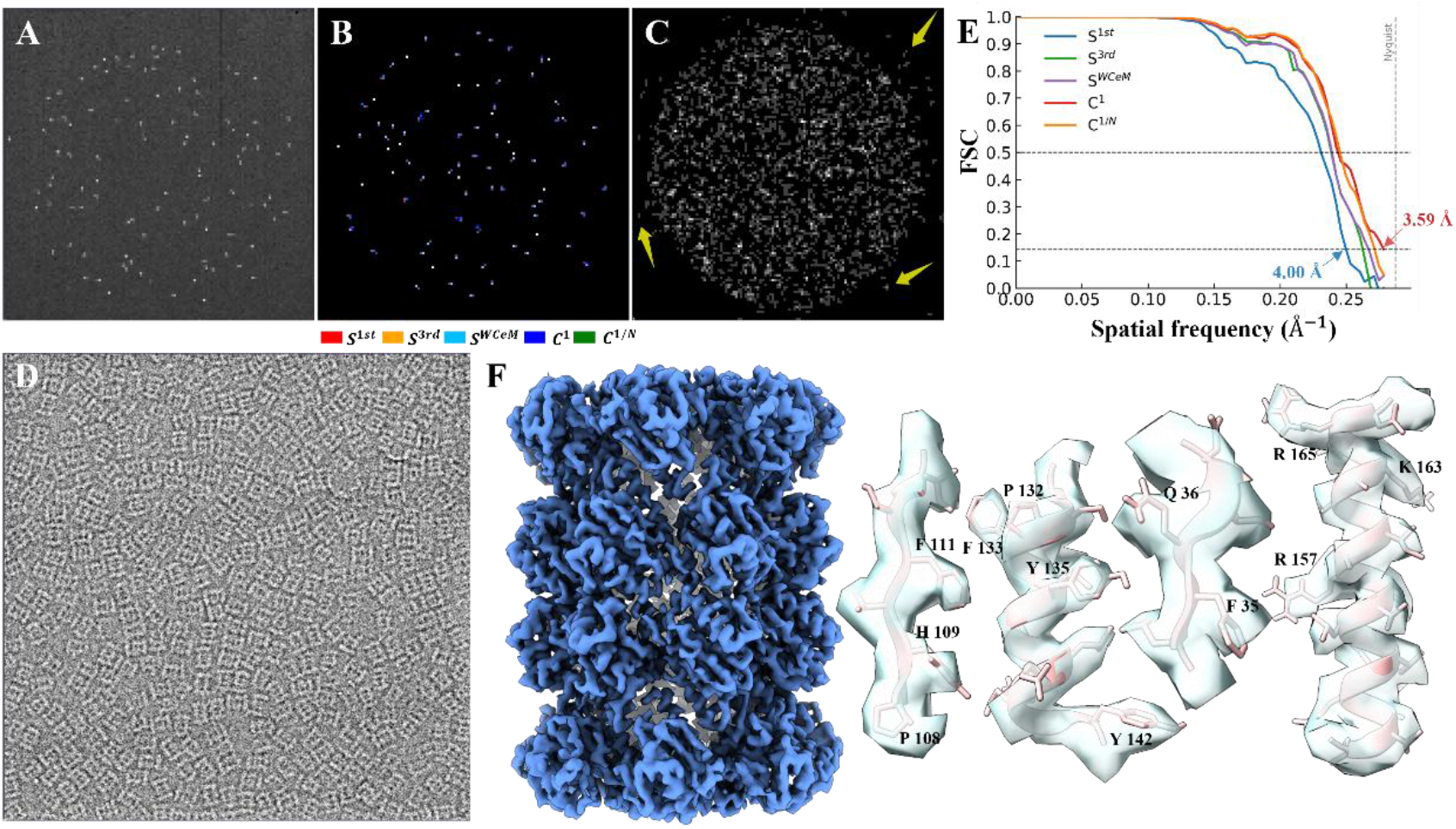
Electron-counting strategies and corresponding SPA results. (**A**) Representative raw detector frame of a CBED image, acquired at a convergence semi-angle of 5.10 mrad. The vertical black line denotes a defective pixel column in the detector. (**B**) Composite visualization of processed results (**fig. S2**) demonstrating the application of multiple counting strategies to panel a), with outcomes from five distinct strategies displayed in separate color channels. (**C**) Integrated sum of 20 detector frames processed using counting strategy *C*^1^. (**D**) Typical ptychographic micrograph of T20S. (**E**) FSC curves of SPA reconstructions corresponding to different counting strategies. (**F**) Density map of T20S reconstructed at 3.59 Å resolution. Several representative densities of the T20S β-subunit within the map are displayed.

SPA reconstructions using different counting strategies all reach near-atomic resolution better than 4 Å, and exhibit significant resolution differences. The maximum resolution difference can reach 0.41 Å. Ranking by FSC curves (**Fig. 2E**), the strategy *S*^1*st*^ gets the lowest resolution, agrees with the position residuals predicted by simulation (**fig. S1C**). Interestingly, the strategy *C*^1^ that sets all pixels in the cluster to 1 electron achieves the best resolution, with the highest resolution of 3.59 Å reported by CryoSPARC. Most side chains are clearly distinguishable in the density map (**Fig. 2F**). This result may suggest that the charge-sharing effect causes the pixel values within clusters to be nearly randomly distributed, and inferring electron incident positions purely based on intensity distribution has considerable uncertainty. Counting all affected pixels as 1 electron may be more conducive to balancing actual electron incident positions. It is noteworthy that the strategy *C*^1/*N*^ that normalizes pixels in a cluster to 1 electron has an FSC that almost overlaps with *C*^1^, just slightly lower at the high-frequency end of the curve. Simply setting a pixel weight of 1/N might not well reflect the weight of such pixel clusters, and more optimal strategies require further research.

In addition to counting strategies, the convergence semi-angle is another key parameter affecting the final reconstruction resolution. We acquired data at a small convergence semi-angle of 3.69 mrad. The electron count per scan position was maintained at approximately 683, corresponding to a total detected electron dose of approximately 7.9 e^−^/Å². This reduced electron count per scan position was implemented to maintain a dose rate within the bright-field disk comparable to that achieved with 5.10 mrad convergence. Because coincidence loss (*10*) represents a critical consideration for electron counting, this configuration was determined to be optimal following extensive experimental trials (data not shown). The ptychographic reconstructed micrographs exhibit superior contrast compared to those obtained at 5.10 mrad (**fig. S4A**). A total of 87,200 selected particles were processed using THUNDER (*15*), ultimately yielding a reconstruction at a resolution of 4.51 Å, which is much lower than that achieved at 5.10 mrad (**fig. S4B** and **C**). Notably, CryoSPARC (*16*) frequently generated unreliable and over-refined high-resolution reconstructions (data not shown), potentially indicating anomalous signals in the high spatial frequency domain of the 3.69-mrad data. Consequently, THUNDER was employed for all analyses and comparisons involving 3.69-mrad data in the following. These findings demonstrate that increasing the convergence semi-angle of the electron probe constitutes a critical factor in attaining near-atomic-resolution in cryo-electron ptychography.

Furthermore, we investigated the contribution of signals located outside the bright-field disk (pointed by yellow arrows in **Fig. 2C**) to SPA, aiming to evaluate the performance of electron counting in detecting these extremely weak signals. The diffraction intensities outside the bright-field disk are remarkably weak (**fig. S5A** and **B**), with corresponding electron counts representing merely approximately 2% of the total dose (**fig. S5C**). Typically, such faint signals would be considered negligible. We carried out comparative ptychographic reconstructions both including and excluding these peripheral signals from the CBED patterns. For both the 3.69-mrad and 5.10-mrad datasets, incorporation of these peripheral signals consistently improved the FSC curves (**fig. S5D**). These findings demonstrate both the excellent performance of electron counting and the practical utility of weak signals outside the bright-field disk for studying frozen hydrated biological specimens.

Taken together, the utilization of electron counting and a large convergence semi-angle constitutes two pivotal factors for achieving near-atomic-resolution ptychographic SPA.

### Ptychographic phase retrieval under low dose

Extending electron ptychography to broader radiation-sensitive applications, especially cryoET, requires ptychographic algorithms to converge robustly under extremely low dose conditions. The typical electron dose per tilt angle in cryoET is 2-3 e^−^/Å², far below the cumulative dose of 20-60 e^−^/Å² in SPA. The adoption of electron counting may provide an opportunity for ptychography at such low dose.

To assess the performance of low-dose ptychographic reconstruction, we generated frame-grouped datasets from the CBED data through electron counting using the following procedure. The 20 detector frames of CBED recorded at each scan position were grouped according to specified frame counts and subsequently summed to produce multiple CBED sets. Each CBED set then underwent independent ptychographic reconstruction to generate corresponding micrograph frames. By integrating micrograph frames from all groupings, we obtained a final composite micrograph suitable for SPA. We examined multiple grouping configurations, specifically testing group sizes of 1, 5, and 10 detector frames, corresponding to distinct ptychography reconstruction dose levels. Importantly, the micrographs produced with varying group sizes maintained consistent total dose, with differences solely attributable to the ptychographic reconstruction performance at different dose levels.

Considering that the image contrast may affect ptychographic reconstruction, we carried out the above experiments separately for 5.10-mrad and 3.69-mrad datasets. For the 5.10-mrad datasets, the group sizes of 1, 5 and 10 detector frames correspond to doses of approximately 1.0, 5.1 and 10.1 e^−^/Å², respectively, used in the ptychographic reconstruction (**Fig. 3A**). Using a micrograph reconstructed with all 20 detector frames at once as the reference, the Fourier ring correlations (FRC) were calculated with the corresponding frame-grouped micrographs to evaluate the influences of reduced dose in ptychographic reconstruction at the micrograph level (**Fig. 3A**). Micrographs with 10-frame and 5-frame groups both ensure FRC higher than 0.9. The FRC of the 1-frame group micrograph shows a slightly noticeable decay with the increasing spatial frequency, but still maintains above 0.74 at Nyquist frequency. Further SPA analysis demonstrates nearly overlapping FSCs for most THUNDER reconstructions obtained using frame-grouped micrographs and reference micrographs (**Fig. 3B**), with only minor deviations observed near the tail regions of the FSCs. The FSC curve from the 1-frame dataset shows less complete overlap compared to others, yet CryoSPARC reports a resolution of 3.65 Å (purple curve in **Fig. 3B**), closely approaching the resolution achieved with the full 20-frame dataset (red curve in **Fig. 2E**). Additional testing with the 3.69 mrad dataset yielded consistent findings (**Fig. 3C** and **D**). These results collectively demonstrate that while phase retrieval at doses as low as 1.0 e/Å² or even 0.4 e/Å² is not a completely lossless process, it can nevertheless preserve high-quality structural information.

**Fig. 3.**
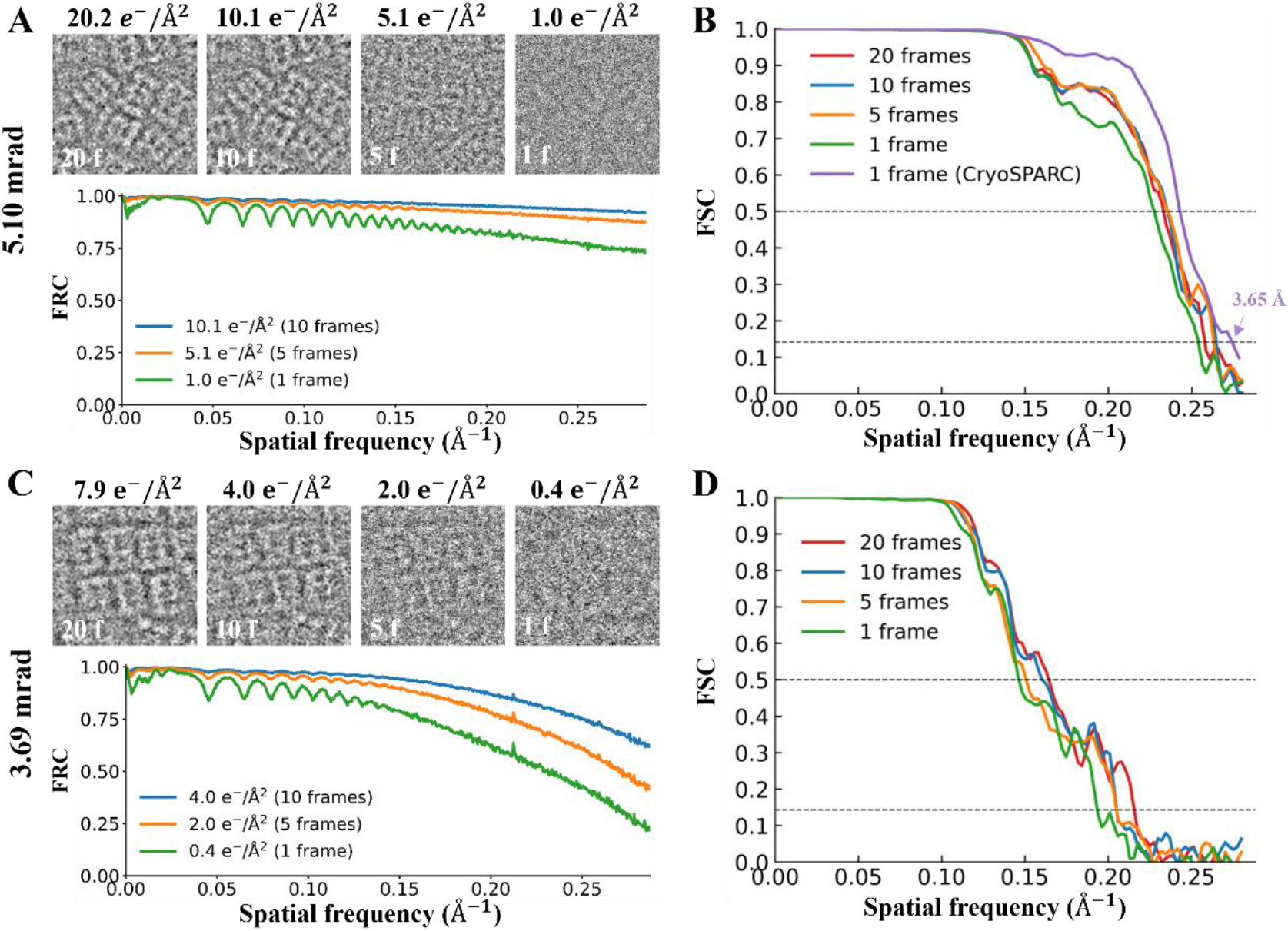
Ptychography under different dose levels. (**A**) For 5.10-mrad data, ptychographic reconstructions of micrograph frames obtained from 20, 10, 5, and 1 grouped detector frames (top panel), along with FRC curves calculated between these micrograph frames (10, 5, and 1 frames) and the reference frame (20 frames). The corresponding doses for each micrograph frame are labeled at the top. (**B**) FSC curves of the 5.10-mrad data calculated using different frame grouping schemes. (**C**) and (**D**) Results from 3.69-mrad data, obtained by applying the same analytical methods as in panels (A) and (B), respectively.

### Low-dose contrast in cryo-electron ptychography

Image contrast is a key metric in cryoEM, significantly impacting downstream processes like 3D reconstruction and structural analysis. In cryo-electron ptychography, the primary factors governing contrast are the electron dose and convergence semi-angle. To evaluate the potential benefits of cryo-electron ptychography in the contrast, we compared it with conventional phase-contrast imaging, specifically under ultra-low-dose conditions of 3 and 6 e^−^/Å² with convergence semi-angles of 3.69 and 5.10 mrad, respectively.

We first examined micrographs of the untilted T20S specimen. The conventional cryoEM micrographs were obtained using defocus values of 2.5 μm and 1.3 μm for comparative analysis (**Fig. 4A**). The conventional micrographs exhibited nearly indiscernible contrast of T20S particles under both low-dose conditions of 3 and 6 e/Å². By contrast, the ptychographic micrographs distinctly revealed identifiable particles, while the particles in the 5.10-mrad micrographs demonstrated reduced contrast compared to those in the 3.69-mrad micrographs. Moreover, for specimens tilted at 10°, 20°, and 30°, the particle contrast became virtually undetectable and progressively diminished with increasing tilt angle (**Fig. 4B**) under phase contrast imaging around 2.5 μm defocus, a condition routinely employed for high-resolution cryoET approaches such as sub-tomogram averaging. It should be noted that the dose utilized in this experiment was 6 e/Å², substantially higher than that typically used in a cryoET tilt angle. In contrast, T20S particles remained clearly discernible in the ptychographic micrograph even at a 30° tilt angle.

**Fig. 4.**
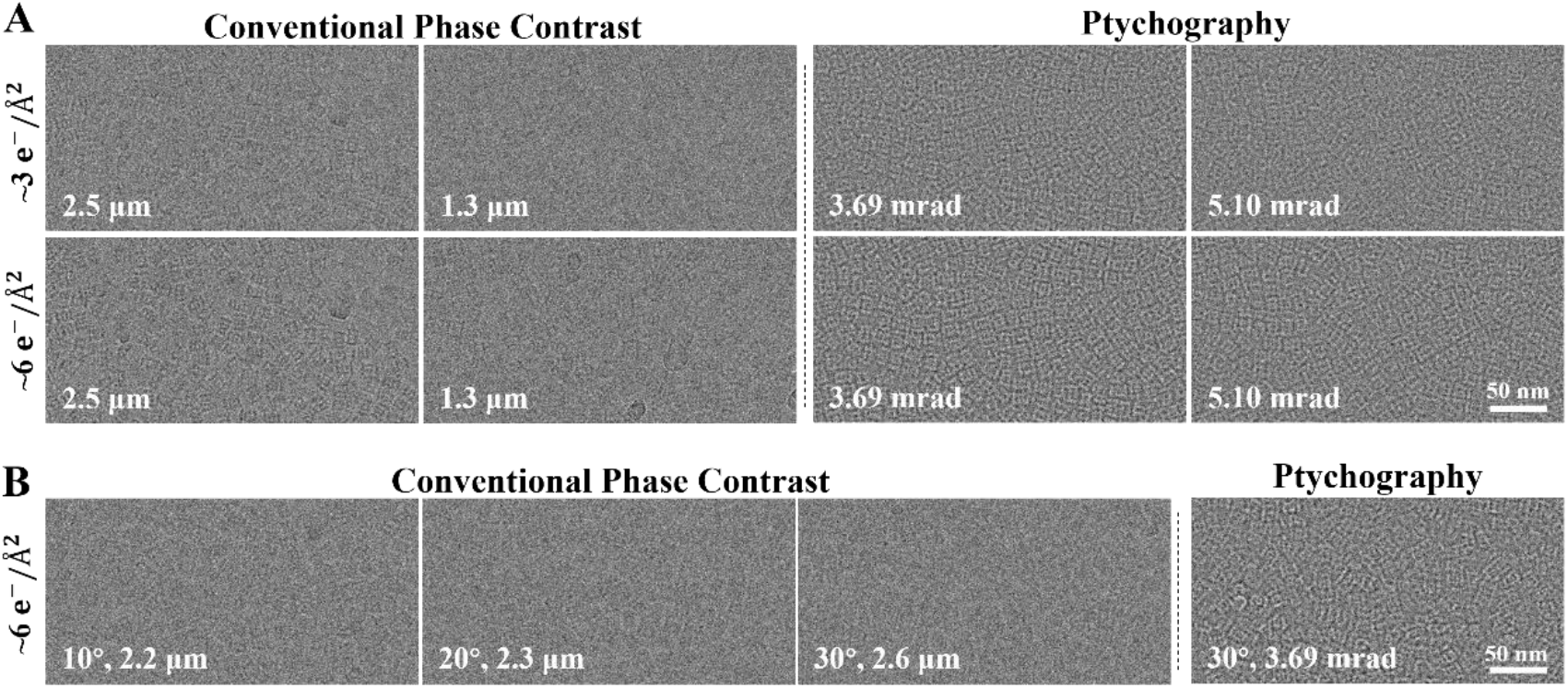
Comparisons of ptychography and conventional phase-contrast imaging under low-dose conditions. All images were binned to a pixel size of 5.22 Å, displayed using a ±3σ grayscale range, and cropped for consistent figure presentation. (**A**) Representative micrographs obtained at dose levels of approximately 3 e^−^/Å² (upper panel) and 6 e^−^/Å² (lower panel), respectively. Conventional phase-contrast imaging employed defocus values of 2.5 μm and 1.3 μm, while ptychography utilized convergence semi-angles of 3.69 mrad and 5.10 mrad accordingly. (**B**) Representative micrographs of tilted specimens acquired at approximately 6 e^−^/Å² dose. Conventional phase-contrast imaging maintained defocus values near 2.5 μm for specimen tilts of 10°, 20°, and 30°, whereas ptychography employed a 3.69 mrad convergence semi-angle for the 30° tilted specimen.

These results, along with the low-dose phase retrieval enabled by electron counting, demonstrate that cryo-electron ptychography offers superior performance for cryoET. Moreover, the convergence semi-angle can be appropriately adjusted based on the desired resolution to optimize image contrast.

### Beam-induced motion under single- and multiple-scan acquisition

Dose fractionation has enabled high-resolution breakthroughs in conventional SPA (*11*). Recent STEM studies on non-biological specimens indicate that multiple-scan strategies can be used to mitigate beam-induced motion (*17*). However, whether the STEM-based dose fractionation is effective or necessary on frozen hydrated biological specimens is to be verified. The scan synchronization control module of EMPIX supports multiple repeated scans of the same sample region and corresponding data acquisition. Utilizing this capability, we separately acquired datasets through single-scan and multiple-scan acquisitions.

In multiple-scan acquisitions, we repeatedly scanned each sample region (350 × 350 scan grid, 3.69 mrad convergence semi-angle) for 4, 5, and 8 times while maintaining the same total dose of 14.8 e^−^/Å². The data from each scan were independently reconstructed using ptychography, generating a multi-frame movie stack for each scanned region. Beam-induced motion was observed in the movie stacks, exhibiting characteristics similar to conventional cryoEM imaging, including rapid global motion during the initial 1-2 frames (**Fig. 5A**) and non-uniform local motion (**Fig. 5B** and **Movies S1**). Various motion patterns were observed (**fig. S6** and **Movies S2-3**), with all movies showing displacements typically ranging approximately from 5 to 90 Å for global motion and up to approximately 20 Å for local motion (**fig. S7**). Following motion correction using MotionCor3, these motion artifacts were effectively eliminated (**Fig. 5C**).

**Fig. 5.**
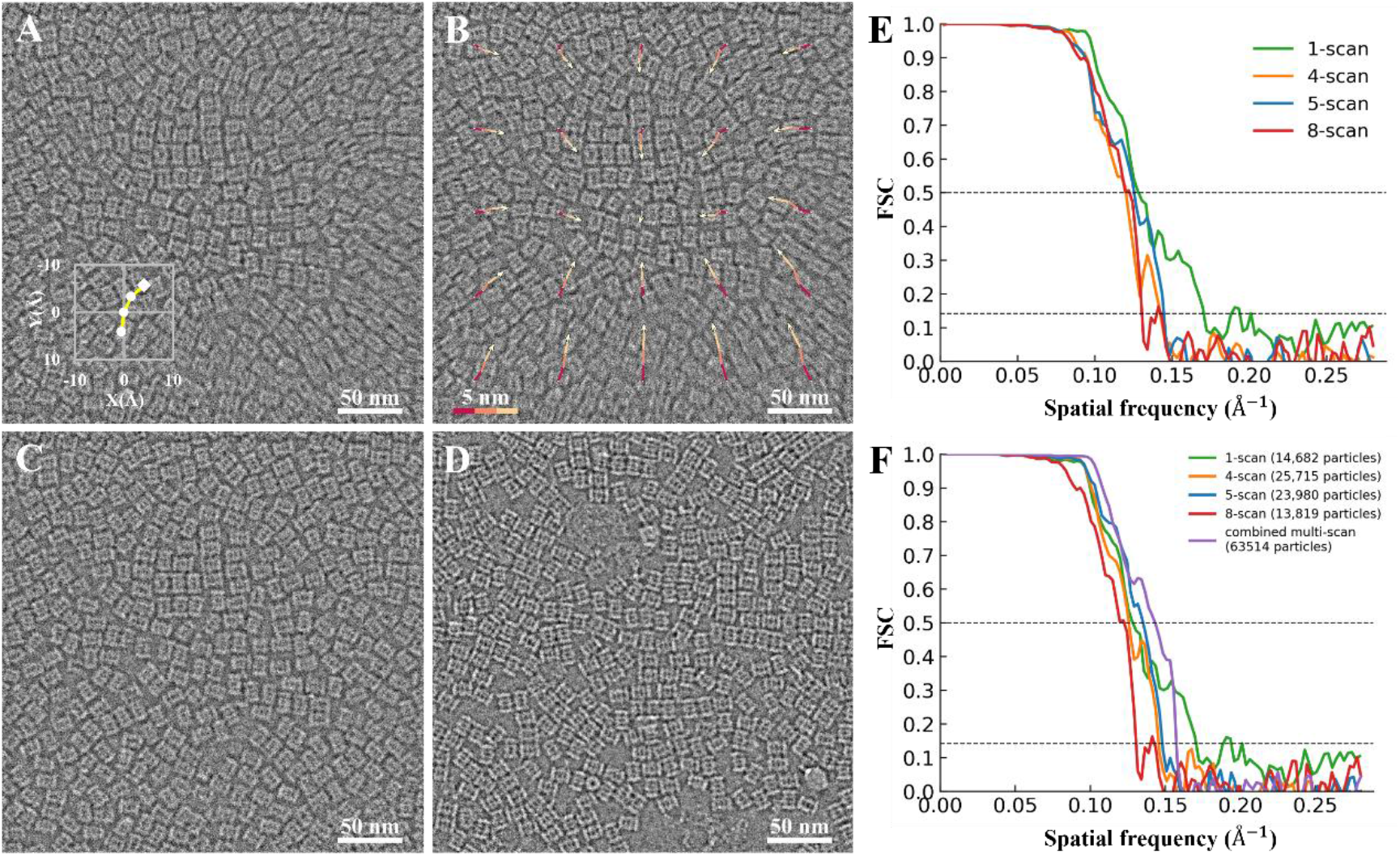
Comparisons of beam-induced motion under multi-scan versus single-scan conditions. A representative micrograph acquired through a multi-scan approach illustrates the motion patterns. (**A**) The global motion observed in the uncorrected micrograph. (**B**) The heterogeneous local motion patterns are clearly discernible in the globally-corrected micrograph, with color-coded traces indicating the measured patch drift trajectories. The trajectories are magnified tenfold, as indicated by the scale bar (bottom left). (**C**) Fully motion-corrected micrograph processed using MotionCor3. (**D**) Representative micrograph obtained via single-scan acquisition. (**E**) FSC curves derived from equivalent particle counts in both single-scan and motion-corrected multi-scan datasets. (**F**) FSC curves computed using complete particle sets from single-scan and motion-corrected multi-scan datasets, including the composite FSC curve generated from all combined multi-scan datasets.

Surprisingly, micrographs obtained through single-scan acquisition exhibit no discernible beam-induced motion (**Fig. 5D**). This observation indicates that beam-induced motion may be absent in single-scan acquisition, or at least is not visibly apparent in the micrographs. To further investigate this unexpected discrepancy in motion characteristics, we performed ptychographic single-particle analysis (SPA) on both the single-scan dataset and the three multi-scan movie datasets. First, we randomly selected 13,819 particles from the single-scan dataset and the motion-corrected multi-scan datasets, respectively, and performed 3D reconstructions using THUNDER. FSC analysis (**Fig. 5E**) reveals that the single-scan dataset achieves higher resolution compared to all multi-scan datasets. Furthermore, within the multi-scan datasets, increasing the number of scans for a single region while maintaining the total dose results in progressive degradation in FSC. Second, we subjected all particles from the four datasets to SPA. The single-scan dataset exhibited only a slightly lower FSC, likely attributable to the relatively fewer particles, yet demonstrated the highest FSC values at the high-resolution tail despite containing nearly the fewest particles (**Fig. 5F**). Notably, even after combining all 63,514 particles from the three multi-scan datasets, the merged reconstruction’s FSC remained inferior in the high-resolution range compared to the single-scan dataset containing merely 14,682 particles.

These results may indicate that during the first scanning round or in a single scan, the probe traverses a fresh specimen where the factors inducing motion have not yet accumulated. Following the first scan, accumulated charging or deformation effects lead to increased motion in subsequent scanning rounds. Notably, even after comprehensive motion correction, the 3D reconstruction quality remains inferior to that obtained from single scans. This observation indicates that current motion correction cannot fully compensate for these accumulated effects, which can be more effectively mitigated through the use of single-scan acquisition. It should be noted that while the single-scan approach avoids charge accumulation and deformation effects between scan rounds, irradiation region overlap persists between adjacent scan positions. The potential existence of corresponding charging or sample deformation phenomena requires further investigation. These findings offer valuable insights for optimizing data acquisition strategies in cryo-electron ptychography and contribute to a different understanding of beam-induced motion mechanisms in frozen hydrated biological specimens.

## Discussion

We have investigated and designed electron-counting strategies utilizing a hybrid-pixel detector and successfully integrated them with ptychography through the acquisition of extremely low-dose CBED data, achieving near-atomic-resolution SPA of the ∼700 kDa T20S proteasome. The optimized electron-counting strategies facilitate effective capture of weak structural information, including sparse diffraction signals outside the bright-field disk that are conventionally excluded, thereby ensuring critical data quality for near-atomic-resolution reconstruction while satisfying various cryoEM prerequisites. These advantages bring superior dose efficiency, enabling the systematic use of much lower doses (≤20.2 e^−^/Å²) in the present work compared to conventional cryoEM. Intriguingly, we observed that scanning transmission imaging may effectively mitigate beam-induced motion, which constitutes a fundamental limitation in conventional SPA.

Among the many factors enabling near-atomic resolution in this work, camera frame rate is an implicit but crucial parameter. A trend in electron microscopes is to pursue higher electron source brightness, yet under our experimental conditions, excessively high electron beam current instead constitutes the main difficulty: it makes it difficult to effectively reduce the dose during scanning to meet dose control requirements. Unlike conventional cryoEM, which can flexibly adjust the dose by reducing the condenser aperture, ptychography has specific requirements for the convergence angle—in this experiment, increasing the probe convergence semi-angle relies on using a larger second condenser aperture (C2 aperture), and increasing the aperture size directly leads to a sharp increase in electron beam current. Under these constraints, when the scan grid size is fixed, the remaining option to control the irradiation dose is to increase the camera frame rate: the lower the dose target, the higher the frame rate is needed to distribute the electron flux to more scan points and detector frames. The maximum frame rate of the camera we currently use is 50 kfps. If a larger convergence semi-angle is desired to further improve resolution, the frame rate may need to reach hundreds of thousands of frames per second to meet the sufficiently low per-frame irradiation dose requirements. Additionally, high frame rate is also a key factor for improving the efficiency of ptychography data acquisition.

The characteristics and operating modes of EMPIX are essential to implement electron counting in diffraction space. The electron counting is achieved based on analysis of charge distribution patterns among adjacent pixels, comprehensively considering the variation of detected electron deposition energy during single electron events. These functionalities are challenging to realize in direct-counting detectors that perform electron counting at the hardware level, as direct inter-pixel communication for coordinated determination of electron incident positions remains difficult. Consequently, under existing technical constraints, determining counting through offline analysis of full-frame signals may represent a more practical compromise solution. For event-driven cameras (*18*), a noteworthy issue is that a single incident electron usually generates response signals simultaneously in multiple adjacent pixels, especially for detectors using smaller pixels. If the event-driven mechanism independently recognizes these multi-pixel responses as multiple electron events, it will heavily occupy bandwidth resources, thereby significantly reducing the number of electrons that the camera can effectively detect per unit time. These may be important factors to consider in the design and implementation of newer detectors in the future.

It should be particularly noted that our recently published work on sampling mismatch has demonstrated that accurate determination of scan step size is another key factor affecting the final SPA reconstruction (*19*). The scan step size used in this study was further corrected based on the newly calculated density map at 3.80 Å resolution. Although the change was only ∼1%, it still shows a clearly visible difference in the FSC curve (**fig. S8A**), with resolution improving from approximately 3.80 Å to 3.59 Å (**fig. S8B**). All results in this work are recalculated based on this updated scan step size. Additionally, all SPA analyses were processed using both THUNDER and CryoSPARC to robustly validate the FSC results (**fig. S9**).

In summary, this study shows that combining cryo-electron ptychography with electron counting and two critical imaging parameters (increased convergence semi-angle and precise scan step size) is essential for enhancing resolution from sub-nanometer to near-atomic levels in frozen hydrated biological specimens. Achieving near-atomic resolution for hundreds kDa protein complexes confirms ptychography’s practical utility in cryoEM, meeting most structural study needs while offering advantages in contrast and beam-induced motion reduction. While we have demonstrated a preliminary application, many imaging capabilities unique to ptychographic computational imaging remain unexplored. These may provide a broader toolkit for complex biological structure characterization, potentially establishing ptychography as a standard cryoEM imaging mode—complementing or even surpassing phase-contrast imaging.

## Data availability

The representative ptychographic datasets and all reconstructed density maps analyzed in this study are available through Zenodo (DOI: 10.5281/zenodo.20758362). The complete ptychographic datasets with raw data, approximately 250 TB in size, will be provided upon request. The T20S proteasome model (PDB:1PMA) was utilized for this study.

## Code availability

The program for ptychographic analysis is the published packaged py4DSTEM (*20*). The executable program for electron counting is available at https://github.com/narutowang/empixCounting-release. The scripts for calibrating the sampling mismatch are available at https://github.com/li-ty22/Sampling-Mismatch-and-Correction-for-Ptychographic-Single-Particle-Analysis.

## Acknowledgements

This work was supported by funds from the National Key Research and Development Program (2024YFA1307302 for X.L. and Z.D.), the National Natural Science Foundation of China (32430056 and 32241023 for X.L.), Beijing Frontier Research Center for Biological Structure, Shenzhen Medical Academy of Research and Translation (SMART) and Tsinghua-Peking Joint Center for Life Sciences. We acknowledge Prof. Binghui Ge from Anhui University for the discussion of 4D-STEM and providing instrument support in the early stage, Dr. Fan Yang for technical assistance with electron microscope operation, Jinying Ma, Ziying Zhang, and Qiaoyang Ma for assistance with sample preparation, and Tianyuan Li for helpful discussions. We acknowledge the Tsinghua University Branch of China National Center for Protein Sciences Beijing for providing facility support in computing and cryoEM instruments.

## Author contributions

X.L. and Z.D. conceived and initiated the project. Z.D. and his team designed and implemented EMPIX. B.S. developed the control software and GUI of EMPIX. S.L., B.S., X.L., and Z.D. jointly developed the electron counting strategies. J.L. and C.T. performed updates and maintenance of the EMPIX camera system. S.L., B.S., and Z.Y. performed data acquisition and processing. Z.W. accelerated the counting code using GPU. J.L. carried out detector simulations and subsequent analysis. X.L., S.L., and J.L. drafted the manuscript. All authors participated in reviewing and revising the manuscript.

## Methods

### Detector simulation

The EMPIX camera is a hybrid-pixel detector characterized by high frame rate and large dynamic range. The camera comprises a 500 μm-thick silicon diode array configured in a 128 × 128 pixel matrix, with a pixel size of 150 μm × 150 μm. The detector employs flip-chip bonding technology to connect to a dedicated readout ASIC (Application Specific Integrated Circuit). Each pixel incorporates a switched integrator, correlated double sampling (CDS) circuit, and Wilkinson-type ADC (Analog-to-Digital Converter), enabling precise charge integration and digitization of detector current signals with 12-bit quantization accuracy. An integrated charge pump circuit extends the dynamic range capability to 24 bits. Through on-chip digitization implementation, the detector achieves a frame rate performance of up to 50 kfps.

The Monte Carlo simulations of the electron detection process in the EMPIX detector were carried out using the open-source simulation platform Allpix2. These simulations encompassed the interaction between incident particles and detector materials, carrier generation and transport, formation of induced current signals, and the response of pixel front-end circuits. The physical detector model comprised a silicon sensor, bonding bumps, and the readout chip. A detector pixel array of 128 × 128 elements was simulated, with only events occurring in the central region being selected for subsequent statistical analysis.

In the simulation, the entire detector is assumed to be in a vacuum environment, with incident particles consisting of 300 keV monoenergetic electrons. The electrons are incident perpendicularly from one side of the detector, with their positions uniformly and randomly distributed within a single pixel. A total of 1000 independent single-electron events were simulated. For each event, the simulation records both the positions and deposited energies of all energy deposition points within the silicon sensitive volume. The energy deposited in silicon is converted to charge based on an average energy requirement of 3.6 eV to produce one electron-hole pair. The sensor is reverse-biased at 160 V, ensuring full depletion of the sensor. Carrier transport employs a linear electric field model, incorporating both drift and diffusion processes. The induced current in each pixel resulting from carrier transport is simulated until complete charge collection is achieved. Electronic noise is introduced using a Gaussian distribution with zero mean, where the RMS value is set to 8000 e^−^ as determined from dark-field measurements. A threshold of 24000 e^−^ is applied, and pixels with amplitudes below this threshold are excluded from subsequent electron counting.

### Electron counting

Electron counting comprises four key steps: threshold determination, connected component identification, electron count estimation, and event position assignment. Prior to the counting procedure, the histogram of pixel value distribution across all detector frames is initially computed, followed by determination of the noise threshold as the valley value between the noise peak and the single-electron response peak corresponding to 300 keV electrons. Under the low-dose conditions employed in this study, single-electron events exhibit sparse distribution across the detector frame (**Fig. 2A**), consequently ensuring that the majority of above-threshold pixel clusters originate from single electron events. For connected component and electron count estimation, we perform 8-neighbor connected component analysis on pixels exceeding the noise threshold. Spatially adjacent pixels are grouped into the same pixel cluster.

Five counting strategies are implemented as follows. (1) Strongest pixel strategy (*S*^1*st*^). The pixel position exhibiting the highest pixel value within the cluster is selected and assigned 1 electron. (2) Third-strongest pixel strategy (*S*^3*rd*^). The pixel position displaying the third-highest pixel value in the cluster is selected and assigned to 1 electron. If the cluster contains fewer than 3 pixels, the pixel with the minimum value is selected instead. (3) Weighted centroid pixel excluding the strongest point (*S^WCeM^*). After eliminating the pixel with the highest pixel value in the cluster, the pixel corresponding to the cluster centroid position - calculated using pixel grayscale values as weights - is selected and assigned 1 electron. (4) All-to-1 strategy (*C*^1^). Each pixel within the cluster is assigned to 1 electron. (5) All-to-1/N strategy (*C*^1/*N*^). Every pixel in the cluster is assigned to 1/N electrons, where N represents the total number of pixels in the cluster.

### Electron microscopy and data collection

A 4 μL drop of purified T20S proteasome solution (∼1.6 mg/mL) was applied to glow-discharged Quantifoil holey carbon grids (Quantifoil Micro Tools GmbH) and subsequently plunge-frozen using a Vitrobot Mark III (FEI).

All ptychographic data were acquired using an EMPIX hybrid-pixel detector mounted on a Thermo Fisher Titan Krios G1 microscope operating at an acceleration voltage of 300 keV. Data were collected in STEM mode, with the convergence semi-angle controlled by the C2 condenser aperture. Two convergence semi-angles were employed, 5.10 mrad and 3.69 mrad, with the probe defocus maintained at approximately 1.5 μm and 2.5 μm, respectively. At each scan position, 20 consecutive detector frames were acquired so that an electron-counting workflow could be applied to each frame, with the cumulative electron dose controlled at approximately 20.2 e^−^/Å² for the 5.10 mrad dataset and 7.9 e^−^/Å² for the 3.69 mrad dataset. For the multi-scan acquisitions, the total number of detector frames per scan position was fixed at 40, corresponding to a total electron dose of 14.8 e^−^/Å² that was distributed equally across the scans: 40 detector frames for a single scan, 10 detector frames per scan for 4 scans, 8 detector frames per scan for 5 scans, and 5 detector frames per scan for 8 scans.

### Ptychography

Ptychographic reconstruction was performed using the open-source software package py4DSTEM. Following electron-counting processing of the detector frames, CBED images were generated by summing the counted detector frames at each scan position. The resulting CBED images were then corrected for bad pixels: detector pixels whose mean value across all scan positions was zero or extremely small (<1×10⁻³) were classified as bad pixels and replaced by the mean of their eight neighboring pixels. In addition, every CBED image acquired with the EMPIX camera was transposed before being fed to the reconstruction algorithm. The initial scan step size of 9.50 Å was obtained by calibrating the STEM scan regions across different magnifications using a TEM standard sample (Cross Grating, S106 of agar) and fitting the calibration curve, and was subsequently refined to 9.41 Å based on the reconstructed T20S density map at 3.80 Å resolution using a previously published step-size calibration method. The initial probe was generated from electron optical parameters, including the convergence semi-angle and defocus value, with the defocus estimated using parallax imaging implemented in py4DSTEM. The object function was initialized as a constant zero electrostatic potential. A single-slice object, single-mode probe reconstruction approach was employed. Joint iterative optimization of the object transmission function and the probe wave function were carried out using a gradient descent algorithm with an update step size of 0.1, and the scan positions were kept fixed throughout the iterations without position correction. All reconstructions were executed on a single NVIDIA L40 GPU. To make full use of the 48 GB on-board memory, the batch size of randomly selected scan positions used for parallel gradient computation was set to 30,720. The total computation time for 30 iterations of a single micrograph was approximately 150 s.

### CryoEM data processing

Motion correction of multi-scan datasets was performed using MotionCor3. Single-particle analysis was carried out using both THUNDER and CryoSPARC (v4.7). All datasets were processed primarily in THUNDER. Particles were first picked from the micrographs by template matching in RELION (*21*), screened through several rounds of 2D classification, and re-extracted using updated coordinates obtained from a preliminary 3D refinement. The re-extracted particle stacks were then imported into THUNDER for 3D refinement using global and local searches only, with CTF refinement disabled. A subsequent masked local refinement was performed using a soft mask whose intensity threshold was chosen to enclose the full protein density while excluding external noise. The 5.10 mrad dataset was additionally processed in CryoSPARC in parallel for cross-validation. In CryoSPARC, the Template Picker function was used for particle picking, and multiple rounds of 2D and 3D classification were employed to eliminate mis-picked particles. The selected particles underwent Homogeneous Refinement followed by Local Refinement, with window_inner_radius and window_outer_radius set to 0.63 and 0.73, respectively.

For the THUNDER processing, a total of 114,050 particles were selected from 462 micrographs for the 5.10 mrad dataset, and 87,200 particles from 372 micrographs for the 3.69 mrad dataset. For the multi-scan datasets acquired at the 3.69 mrad convergence semi-angle, 14,682 particles were selected from 57 single-scan (1-frame) micrographs, 25,715 particles from 103 4-frame micrographs, 23,980 particles from 108 5-frame micrographs, and 13,819 particles from 83 8-frame micrographs. For the CryoSPARC processing, a total of 109,227 particles were selected from 462 micrographs for the 5.10 mrad dataset. When directly comparing single-particle analysis results across datasets, all datasets were randomly subsampled to the minimum particle count. The final resolution was determined using the FSC criterion at 0.143, and all density-map visualizations were generated using ChimeraX.

## Supplementary Figures and Legends

**Fig. S1.**
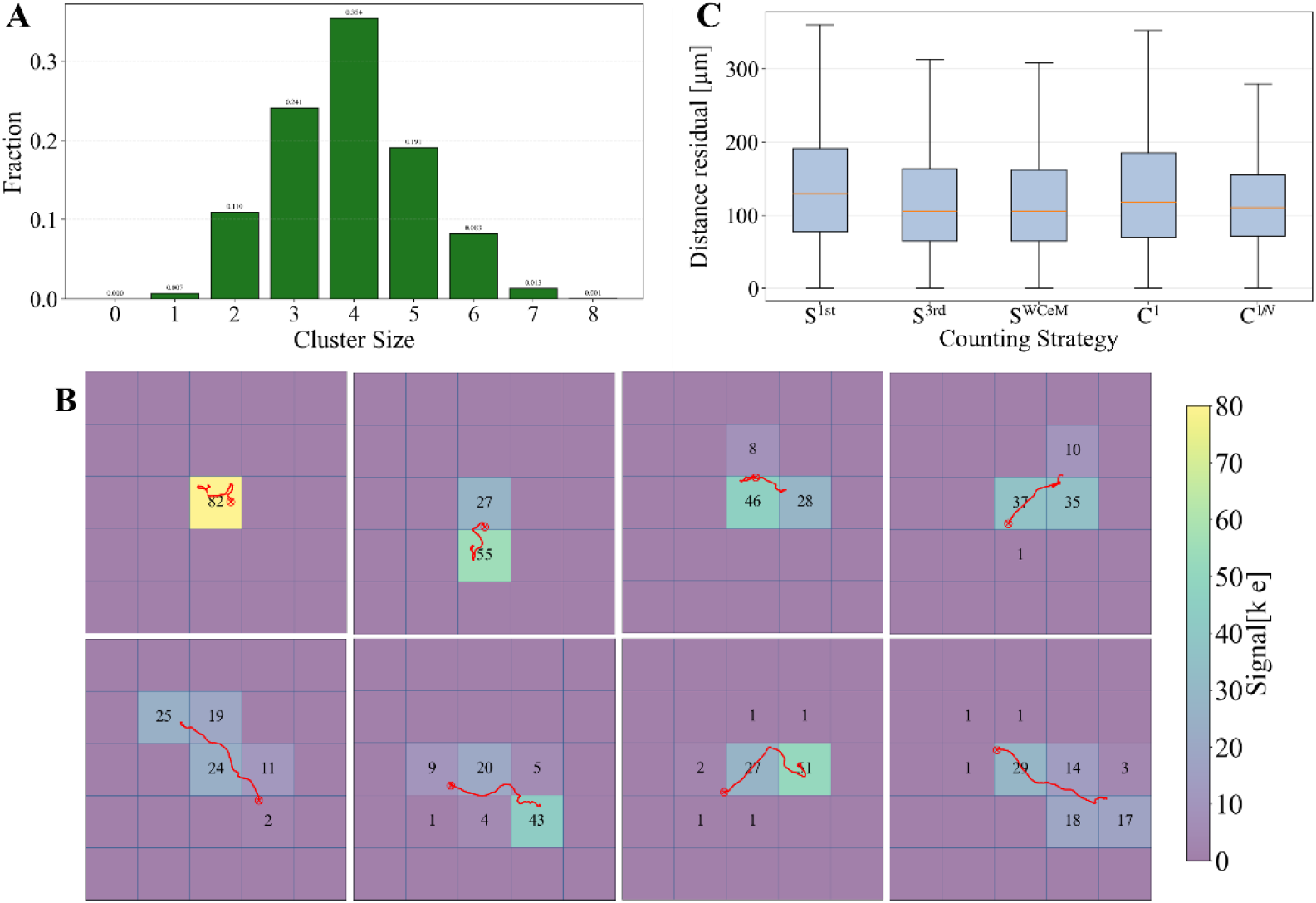
Pixel clusters and counting position residuals for different counting strategies, derived from simulation data. (**A**) Size distribution of pixel clusters impacted by simulated single-electron events without accounting for background subtraction. In contrast to Fig. 1D, this statistic includes all pixels within each cluster without excluding those below the noise threshold. (**B**) Representative example of simulated pixel clusters affected by a single-electron event. (**C**) Counting position residuals corresponding to five distinct counting strategies.

**Fig. S2.**
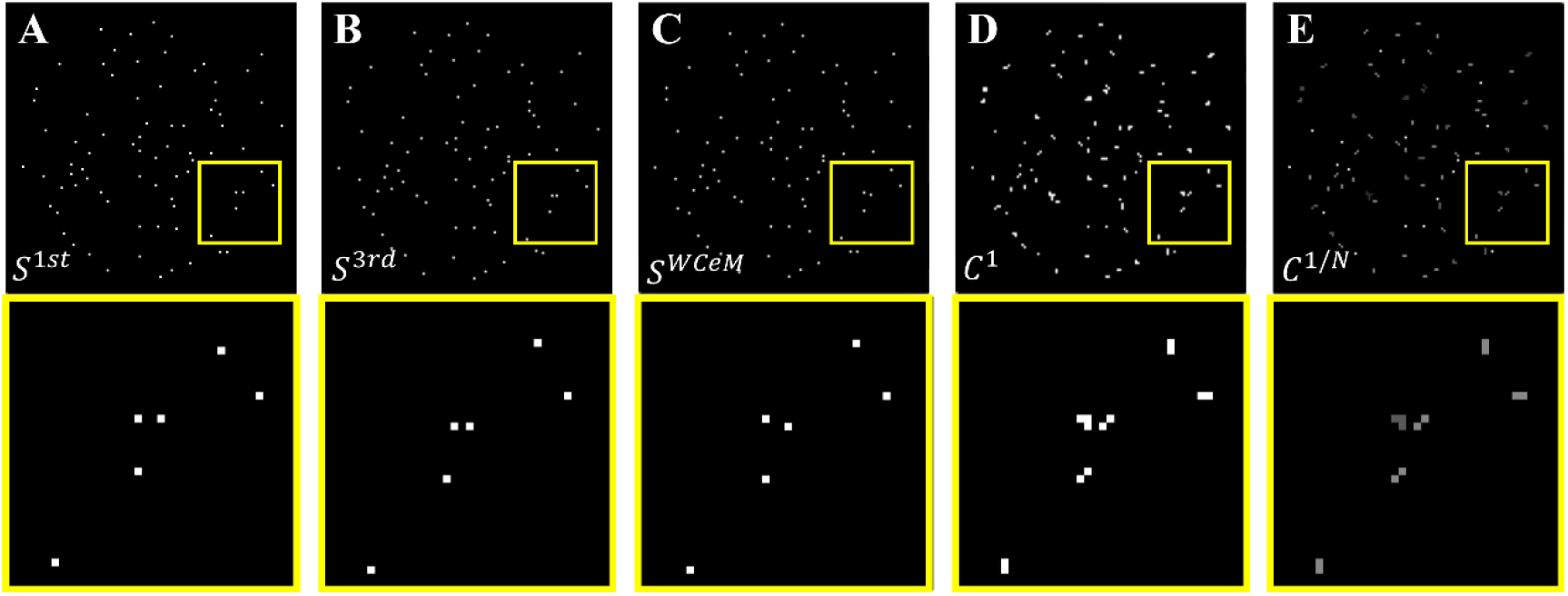
Comparison of electron counting using different counting strategies. The detector frame depicted in Fig. 1A is tested here. (**A**) – (**E**) Counting results obtained by applying the specific strategy indicated in the bottom left corner. For enhanced visualization, a magnified view of the local region marked by the yellow box is displayed below each panel. The analysis reveals that different strategies assign distinct values to identical charge-sharing clusters, resulting in subtle positional variations. The first three strategies exclusively select a single pixel position within the cluster to represent the incident position. In contrast, the latter two strategies incorporate all pixels within the cluster: the *C*^1^ strategy assigns an intensity value of 1 to every pixel in the cluster, whereas the *C*^1/*N*^ strategy demonstrates an inverse relationship between pixel intensity and cluster size, with individual pixel intensity decreasing as the number of pixels in the cluster increases.

**Fig. S3.**
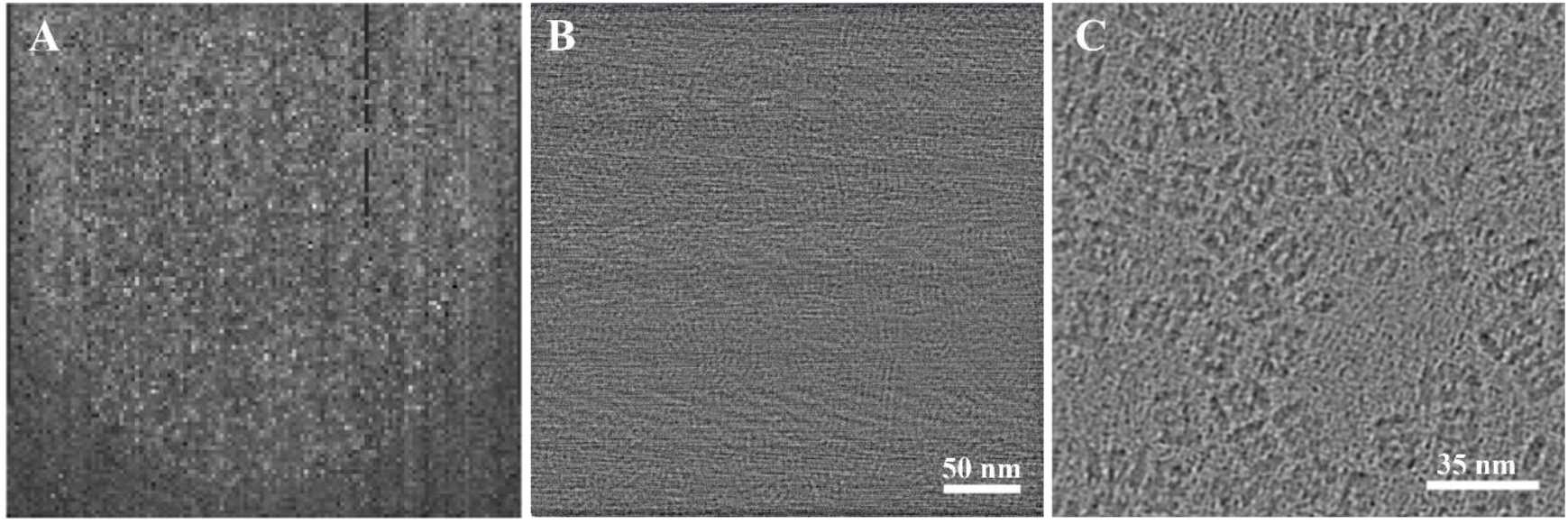
Ptychography reconstruction without electron counting. (**A**) A low-dose CBED image obtained by merging 20 detector frames without electron counting. (**B**) Low-dose ptychographic reconstruction of the CBED dataset containing the image shown in panel (A). The reconstructed image exhibits severe modulation by periodic stripe noise, with poor contrast of the T20S proteasome particles. (**C**) Ptychographic reconstruction of CBED data acquired from a negative-stained T20S proteasome sample under high-dose conditions, where the particles are clearly discernible. This comparative analysis demonstrates that for low-dose imaging of frozen-hydrated biological specimens, electron counting is crucial for enhancing both diffraction data quality and subsequent ptychographic reconstruction quality.

**Fig. S4.**
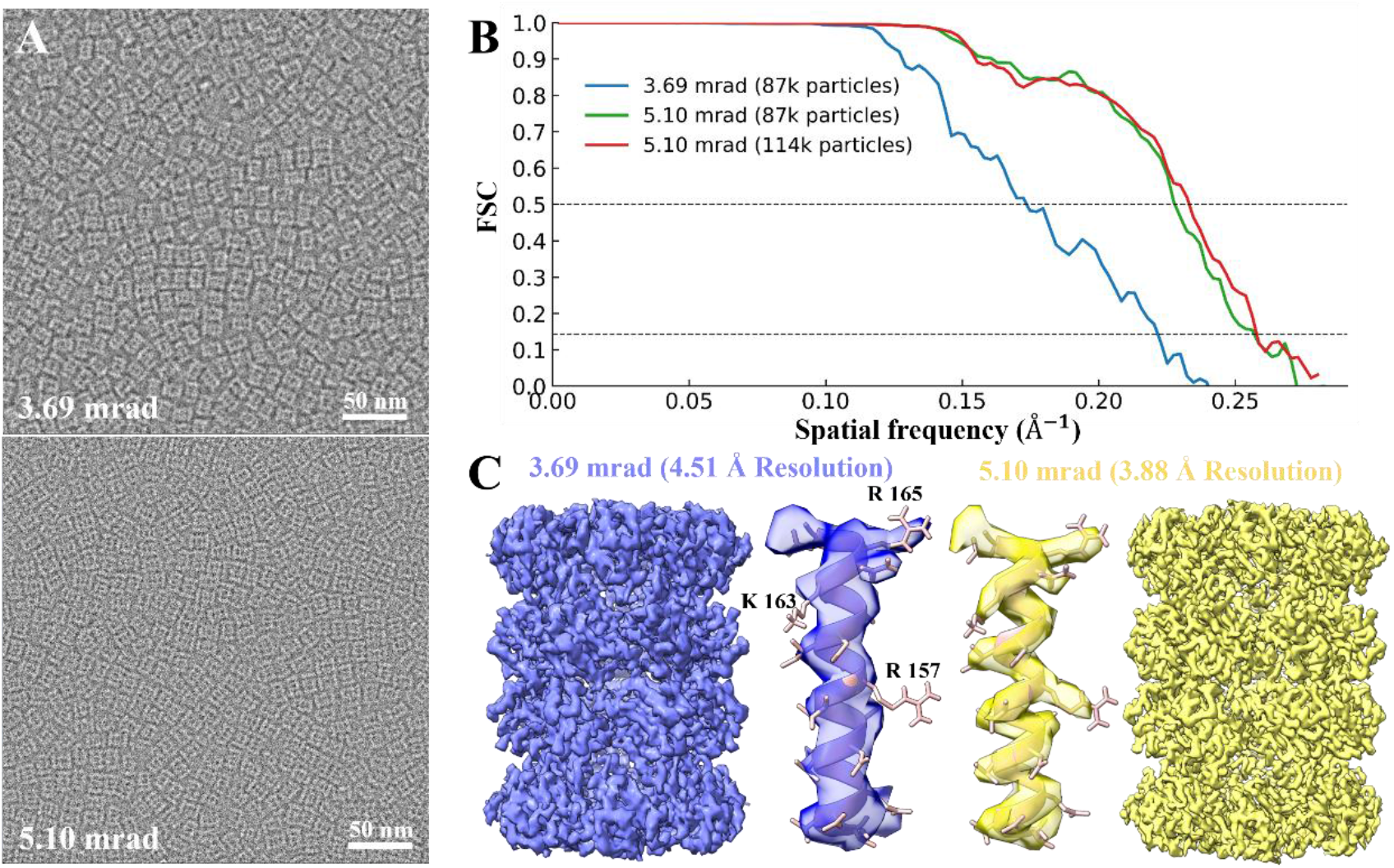
Comparison of cryo-electron ptychography reconstruction results at convergence semi-angles of 3.69 mrad and 5.10 mrad. (**A**) Representative micrographs acquired at the two convergence semi-angles. The smaller 3.69 mrad convergence semi-angle demonstrates superior image contrast compared to the larger 5.10 mrad convergence semi-angle. (**B**) FSC curve comparison between datasets obtained at different convergence semi-angles. The single-particle analysis (SPA) using the 5.10-mrad dataset exhibits much better FSC characteristics and higher resolution. Two FSC curves are calculated for the 5.10-mrad dataset: one utilizing all available particles and another employing a matched particle count equivalent to that of the 3.69-mrad dataset. (**C**) Structural comparison of single-particle analysis density maps reconstructed at both convergence semi-angles. The density map obtained at 5.10 mrad convergence semi-angle displays significantly enhanced side-chain density features relative to the 3.69 mrad reconstruction. These representative side-chain densities are from the T20S β-subunit within the map.

**Fig. S5.**
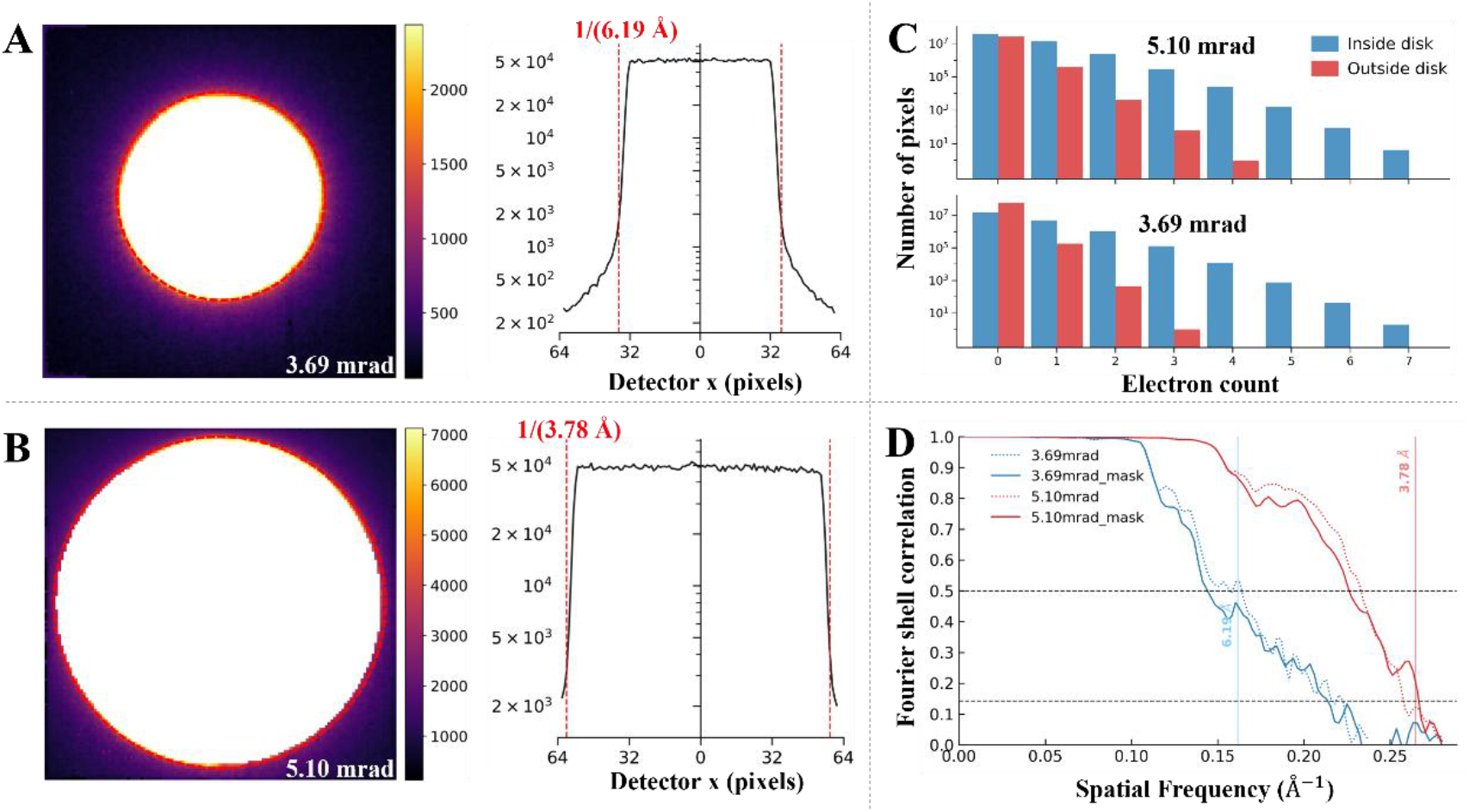
Intensity distribution of weak signals outside the bright-field disk and their influence on 3D reconstruction. All panels below display electron-counted data following bad-pixel correction. (**A**) Averaged CBED image obtained by integrating the 3.69 mrad convergence semi-angle data across all scan positions (left), with the corresponding intensity profile through the center of the bright-field disk (right). (**B**) Averaged CBED image for the 5.10 mrad convergence semi-angle data, presented in the same format as panel (A). In both (A) and (B), a circular mask (denoted by red circles and dashed lines) with radius matching the bright-field disk is applied to estimate weak-signal detection outside. The radius is numerically indicated in red within the profile panel. (**C**) Histogram showing the electron-count distribution for detector pixels located inside versus outside the bright-field disk mask. The external signals constitute approximately 2.45% of total signal intensity for the 3.69-mrad data and 1.88% for the 5.10-mrad data. (**D**) Comparison of FSC curves derived from SPA, contrasting results obtained using only signals within the bright-field disk mask versus full detector signals. Blue and red vertical lines indicate the real-space resolutions corresponding to mask boundaries for the 3.69 mrad and 5.10 mrad datasets, respectively. Under both convergence semi-angles, the FSC curves incorporating full detector signals demonstrate consistently superior performance compared to those utilizing only the masked bright-field disk signals.

**Fig. S6.**
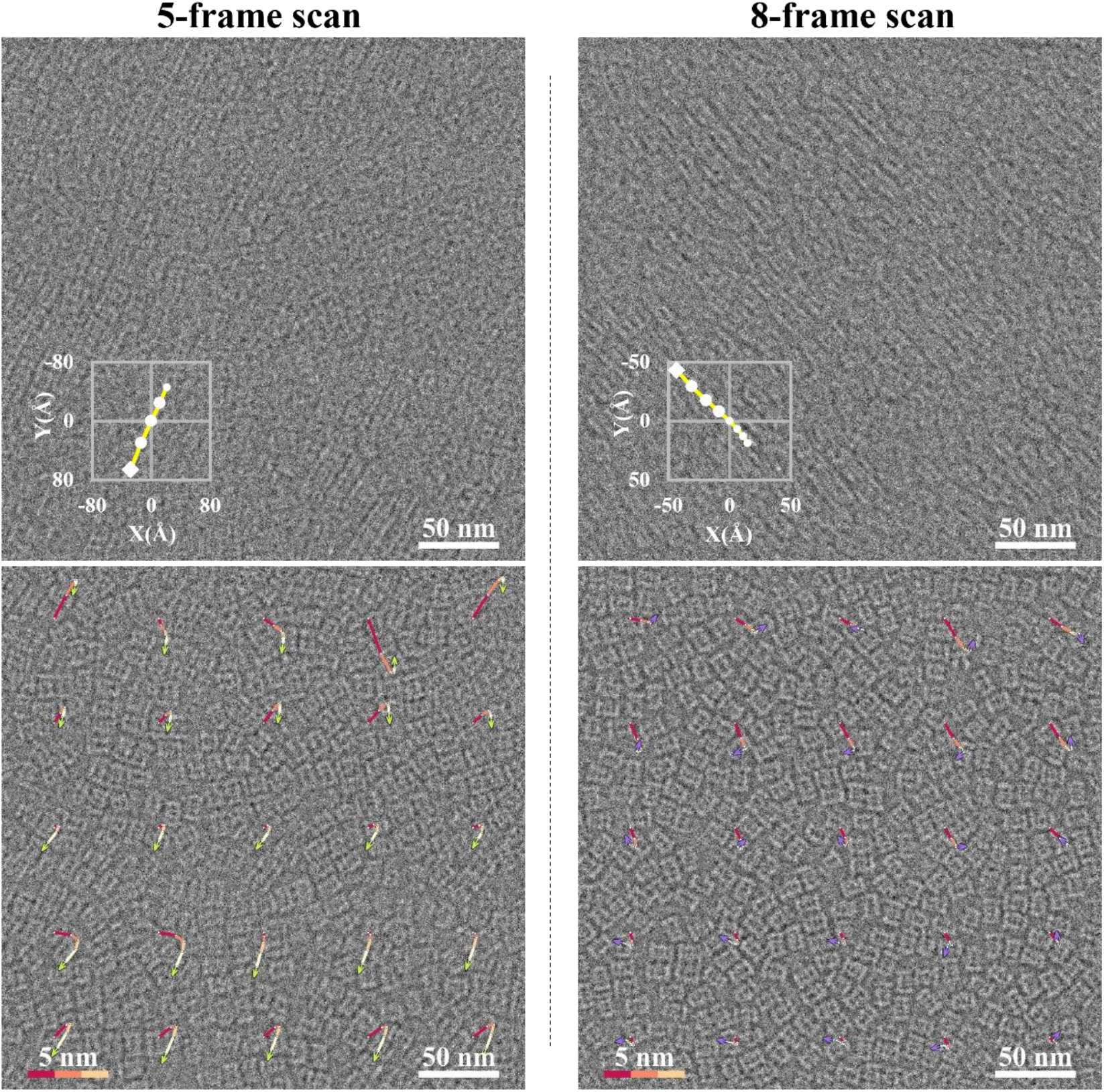
Beam-induced motion patterns observed in multi-scan acquisitions. As an extension to Fig. 5A and **C** (based on 4-scan acquisitions), this figure presents representative motion patterns obtained from 5-scan and 8-scan acquisitions. The corresponding video demonstrations are provided in **Movies S2** and **3**. The upper panels display global drift trajectories, while the lower panels present non-uniform local drift maps, consistent with the format shown in Fig. 5. The trajectories are magnified tenfold, as indicated by the scale bar (bottom left).

**Fig. S7.**
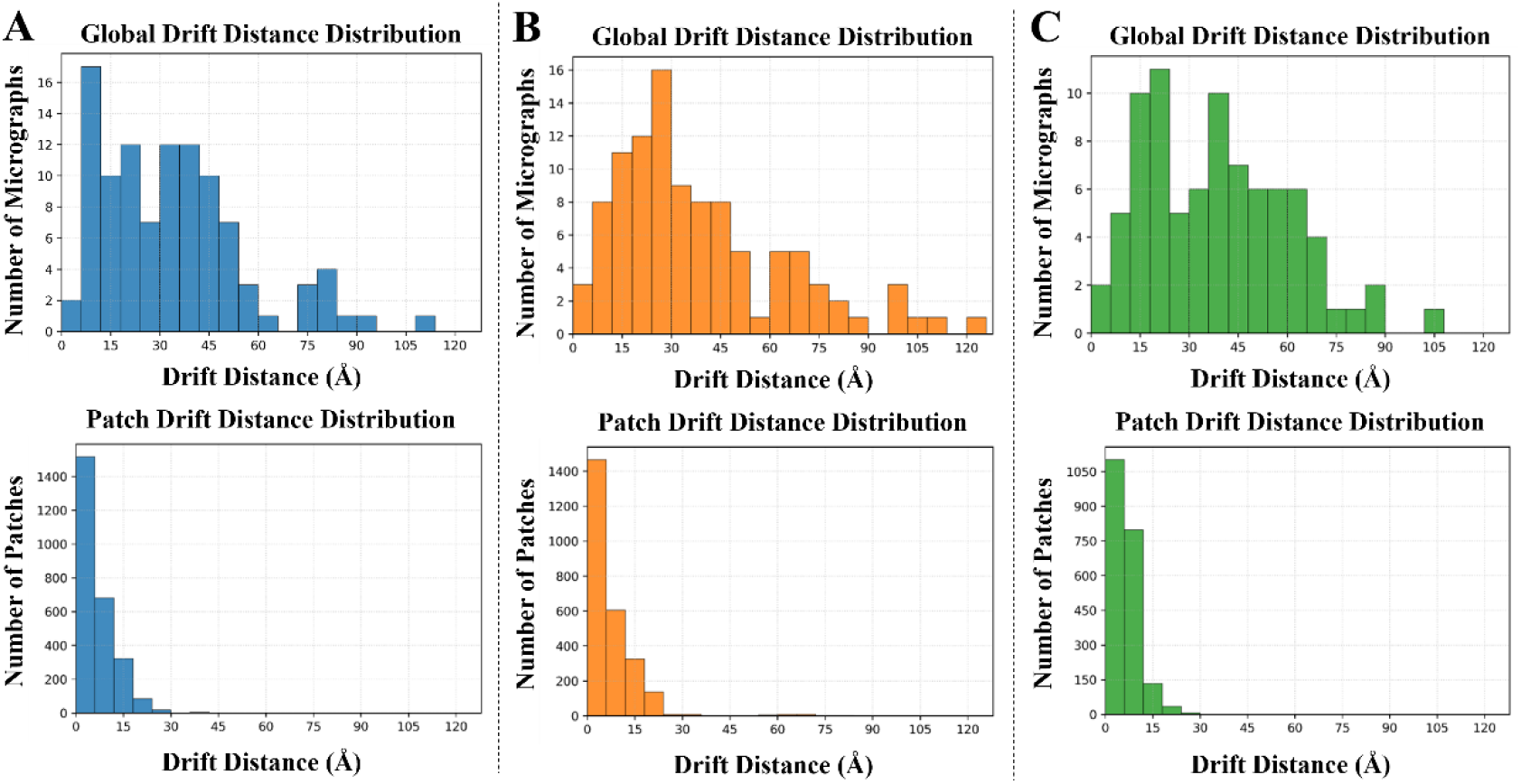
Histograms of global drift and local patch drift distances for datasets acquired with varying numbers of multi-scans. (**A**) – (**C**) Statistical results for 4-scan, 5-scan, and 8-scan acquisitions. The distributions of both global and patch drift demonstrate substantial consistency across different scan counts. Notably, patch drift predominantly remains within 20 Å, with over 50% of cases falling below 6 Å, whereas global drift primarily distributes between 5–90 Å.

**Fig. S8.**
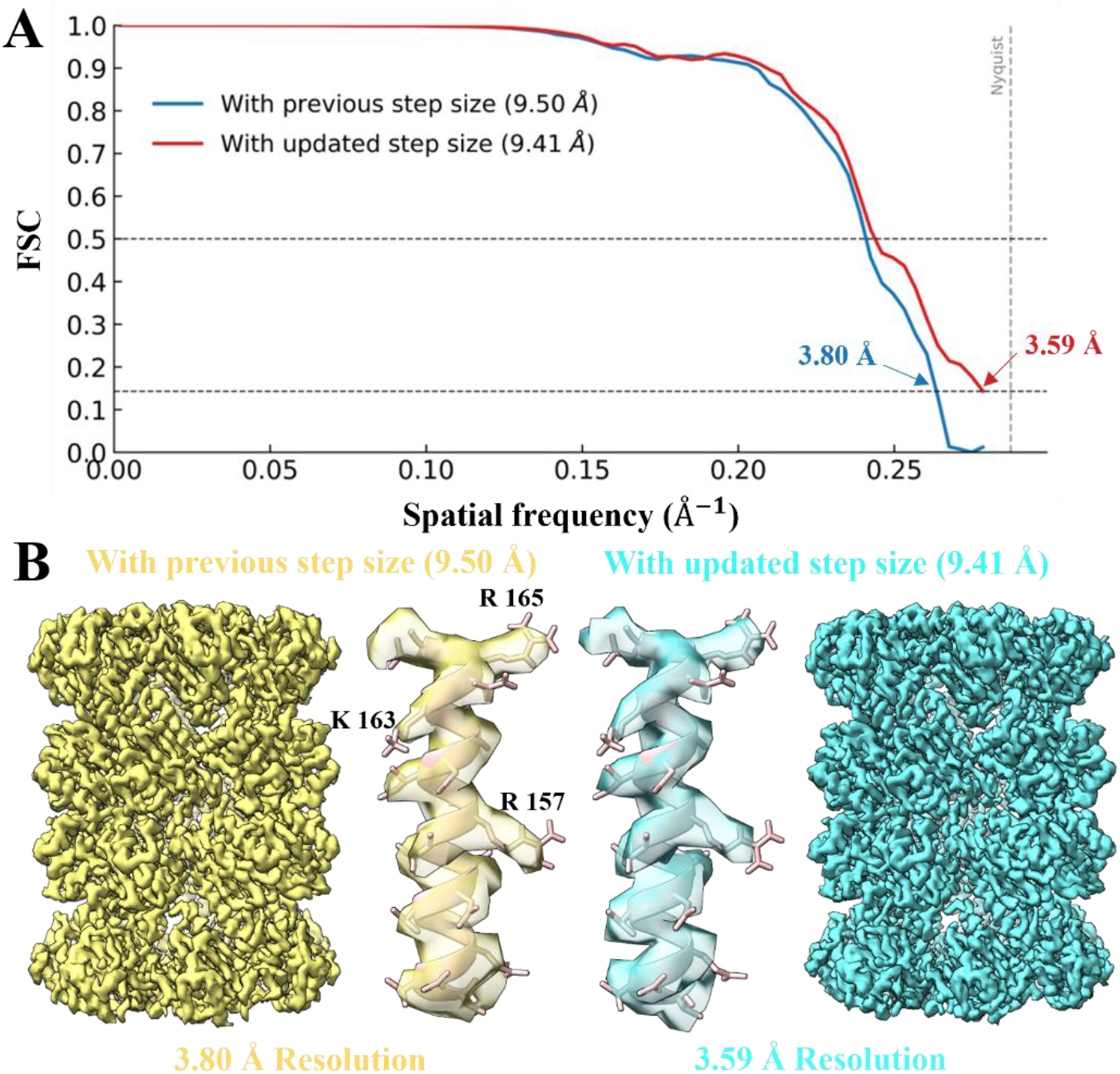
Influence of scan step size on ptychography reconstruction and SPA. (**A**) Comparison of FSC curves from SPA obtained using the original (9.50 Å) and optimized (9.41 Å) scan step sizes. The FSC curve corresponding to the optimized scan step size demonstrates improvement across all spatial frequencies, with the resolution enhancing from approximately 3.80 Å to 3.59 Å. (**B**) Comparison of reconstructed density maps generated with the previous and updated scan step sizes. A representative α-helix density from the T20S β-subunit is displayed for each map. The density map reconstructed using the updated step size exhibits enhanced structural detail. The previous scan step size of 9.50 Å was determined through calibration using a reconstructed density map at ∼6 Å resolution. The optimized scan step size of 9.41 Å was subsequently derived through further refinement based on the initial T20S density map at 3.80 Å resolution. Although the adjustment in step size represents only approximately a 1% change, the FSC curve nevertheless displays measurable improvement, underscoring that precise scan step size calibration constitutes a critical processing parameter for achieving near-atomic resolution in cryo-electron ptychography.

**Fig. S9.**
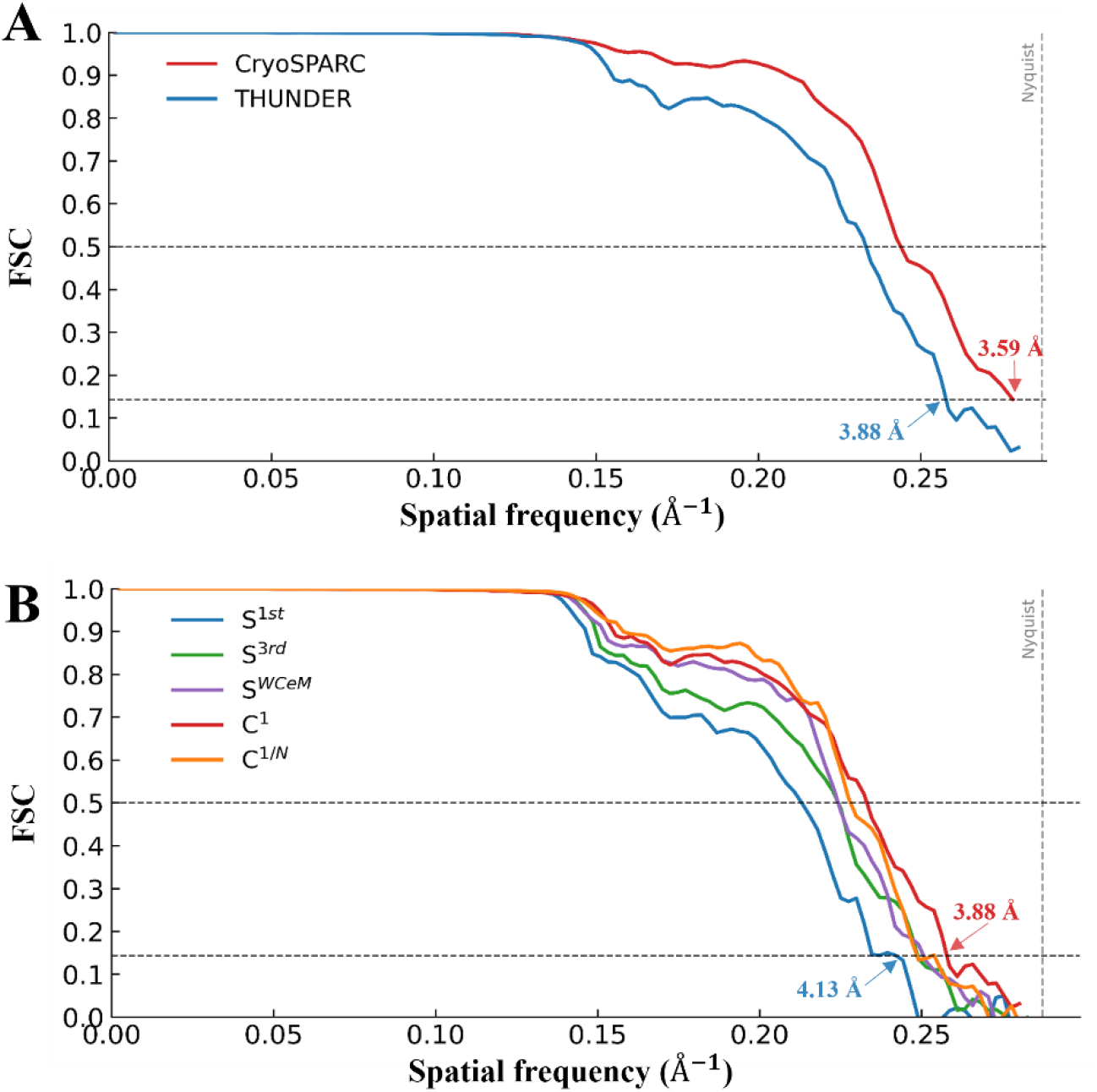
Comparative analysis of THUNDER and CryoSPARC single-particle processing results for the 5.10 mrad convergence semi-angle dataset. (**A**) FSC curves from SPA of the 5.10-mrad dataset processed with THUNDER and CryoSPARC. CryoSPARC demonstrates superior performance in the FSC curve and reports higher resolution (3.59 Å), while the THUNDER reconstruction attains a resolution of 3.88 Å. (**B**) FSC curves from the 5.10 mrad dataset using different counting strategies. Both THUNDER and CryoSPARC analyses yield consistent conclusions, with the *C*^1^ strategy producing the highest resolution and the *S*^1*st*^ strategy resulting in the lowest resolution.

## References and Notes

1. C. Ophus, P. Ercius, M. Sarahan, C. Czarnik, J. Ciston, Recording and Using 4D-STEM Datasets in Materials Science. Microanal 20, 62–63 (2014).

2. Y. Jiang, Z. Chen, Y. Han, P. Deb, H. Gao, S. Xie, P. Purohit, M. W. Tate, J. Park, S. M. Gruner, V. Elser, D. A. Muller, Electron ptychography of 2D materials to deep sub-ångström resolution. Nature 559, 343–349 (2018).

3. Z. Chen, Y. Jiang, Y.-T. Shao, M. E. Holtz, M. Odstrčil, M. Guizar-Sicairos, I. Hanke, S. Ganschow, D. G. Schlom, D. A. Muller, Electron ptychography achieves atomic-resolution limits set by lattice vibrations. Science 372, 826–831 (2021).

4. K. X. Nguyen, Y. Jiang, C.-H. Lee, P. Kharel, Y. Zhang, A. M. van der Zande, P. Y. Huang, Achieving sub-0.5-angstrom–resolution ptychography in an uncorrected electron microscope. Science 383, 865–870 (2024).

5. X. Pei, L. Zhou, C. Huang, M. Boyce, J. S. Kim, E. Liberti, Y. Hu, T. Sasaki, P. D. Nellist, P. Zhang, D. I. Stuart, A. I. Kirkland, P. Wang, Cryogenic electron ptychographic single particle analysis with wide bandwidth information transfer. Nat Commun 14, 3027 (2023).

6. B. Küçükoğlu, I. Mohammed, R. C. Guerrero-Ferreira, S. M. Ribet, G. Varnavides, M. L. Leidl, K. Lau, S. Nazarov, A. Myasnikov, M. Kube, J. Radecke, C. Sachse, K. Müller-Caspary, C. Ophus, H. Stahlberg, Low-dose cryo-electron ptychography of proteins at sub-nanometer resolution. Nat Commun 15, 8062 (2024).

7. P. M. Pelz, W. X. Qiu, R. Bücker, G. Kassier, R. J. D. Miller, Low-dose cryo electron ptychography via non-convex Bayesian optimization. Sci Rep 7, 9883 (2017).

8. L. Zhou, J. Song, J. S. Kim, X. Pei, C. Huang, M. Boyce, L. Mendonça, D. Clare, A. Siebert, C. S. Allen, E. Liberti, D. Stuart, X. Pan, P. D. Nellist, P. Zhang, A. I. Kirkland, P. Wang, Low-dose phase retrieval of biological specimens using cryo-electron ptychography. Nat Commun 11, 2773 (2020).

9. Y. Yu, K. A. Spoth, M. Colletta, K. X. Nguyen, S. E. Zeltmann, X. S. Zhang, M. Paraan, M. Kopylov, C. Dubbeldam, D. Serwas, H. Siems, D. A. Muller, L. F. Kourkoutis, Dose-efficient cryo-electron microscopy for thick samples using tilt-corrected scanning transmission electron microscopy. Nat Methods 22, 2138–2148 (2025).

10. X. Li, S. Q. Zheng, K. Egami, D. A. Agard, Y. Cheng, Influence of electron dose rate on electron counting images recorded with the K2 camera. Journal of Structural Biology 184, 251–260 (2013).

11. X. Li, P. Mooney, S. Zheng, C. R. Booth, M. B. Braunfeld, S. Gubbens, D. A. Agard, Y. Cheng, Electron counting and beam-induced motion correction enable near-atomic-resolution single-particle cryo-EM. Nat Methods 10, 584–590 (2013).

12. A. R. Faruqi, G. McMullan, Direct imaging detectors for electron microscopy. Nuclear Instruments and Methods in Physics Research Section A: Accelerators, Spectrometers, Detectors and Associated Equipment 878, 180–190 (2018).

13. T. Wei, H. Yang, Z. Deng, T. Xue, X. Wang, Design and evaluation of EMPIX2, a 100 kfps, high dynamic range pixel detector readout ASIC for electron microscopy. J. Inst. 18, C12007 (2023).

14. S. Spannagel, K. Wolters, D. Hynds, N. Alipour Tehrani, M. Benoit, D. Dannheim, N. Gauvin, A. Nürnberg, P. Schütze, M. Vicente, Allpix2: A modular simulation framework for silicon detectors. Nuclear Instruments and Methods in Physics Research Section A: Accelerators, Spectrometers, Detectors and Associated Equipment 901, 164–172 (2018).

15. M. Hu, H. Yu, K. Gu, Z. Wang, H. Ruan, K. Wang, S. Ren, B. Li, L. Gan, S. Xu, G. Yang, Y. Shen, X. Li, A particle-filter framework for robust cryo-EM 3D reconstruction. Nat Methods 15, 1083–1089 (2018).

16. A. Punjani, J. L. Rubinstein, D. J. Fleet, M. A. Brubaker, cryoSPARC: algorithms for rapid unsupervised cryo-EM structure determination. Nat Methods 14, 290–296 (2017).

17. D. Jannis, C. Hofer, C. Gao, X. Xie, A. Béché, T. J. Pennycook, J. Verbeeck, Event driven 4D STEM acquisition with a Timepix3 detector: Microsecond dwell time and faster scans for high precision and low dose applications. Ultramicroscopy 233, 113423 (2022).

18. R. Ballabriga, J. A. Alozy, F. N. Bandi, G. Blaj, M. Campbell, P. Christodoulou, V. Coco, A. Dorda, S. Emiliani, K. Heijhoff, E. Heijne, T. Hofmann, J. Kaplon, A. Koukab, I. Kremastiotis, X. Llopart, M. Noy, A. Paterno, M. Piller, J. M. Sallesse, V. Sriskaran, L. Tlustos, M. Van Beuzekom, The Timepix4 analog front-end design: Lessons learnt on fundamental limits to noise and time resolution in highly segmented hybrid pixel detectors. Nuclear Instruments and Methods in Physics Research Section A: Accelerators, Spectrometers, Detectors and Associated Equipment 1045, 167489 (2023).

19. T. Li, S. Li, Z. Yan, Y. Shen, X. Li, Sampling Mismatch and Correction for Ptychographic Single-Particle Analysis. bioRxiv [Preprint] (2026). 10.64898/2026.02.21.707235.

20. B. H. Savitzky, S. E. Zeltmann, L. A. Hughes, H. G. Brown, S. Zhao, P. M. Pelz, T. C. Pekin, E. S. Barnard, J. Donohue, L. Rangel DaCosta, E. Kennedy, Y. Xie, M. T. Janish, M. M. Schneider, P. Herring, C. Gopal, A. Anapolsky, R. Dhall, K. C. Bustillo, P. Ercius, M. C. Scott, J. Ciston, A. M. Minor, C. Ophus, py4DSTEM: A Software Package for Four-Dimensional Scanning Transmission Electron Microscopy Data Analysis. Microsc Microanal 27, 712–743 (2021).

21. S. H. W. Scheres, RELION: Implementation of a Bayesian approach to cryo-EM structure determination. Journal of Structural Biology 180, 519–530 (2012).

