## Supplementary figures and images for "Electron counting enables cryo-electron ptychography for near-atomic-resolution cryo-electron microscopy"

### Movie S1

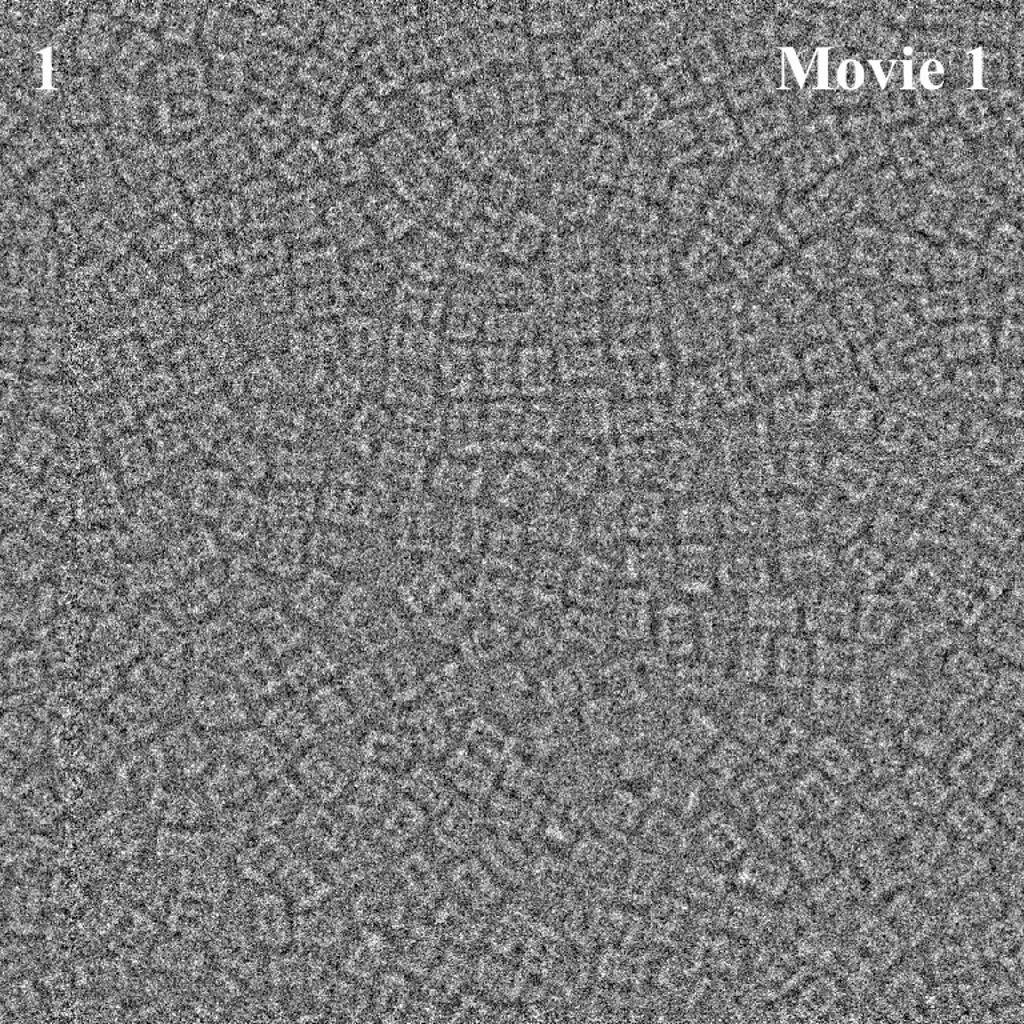

### Movie S2

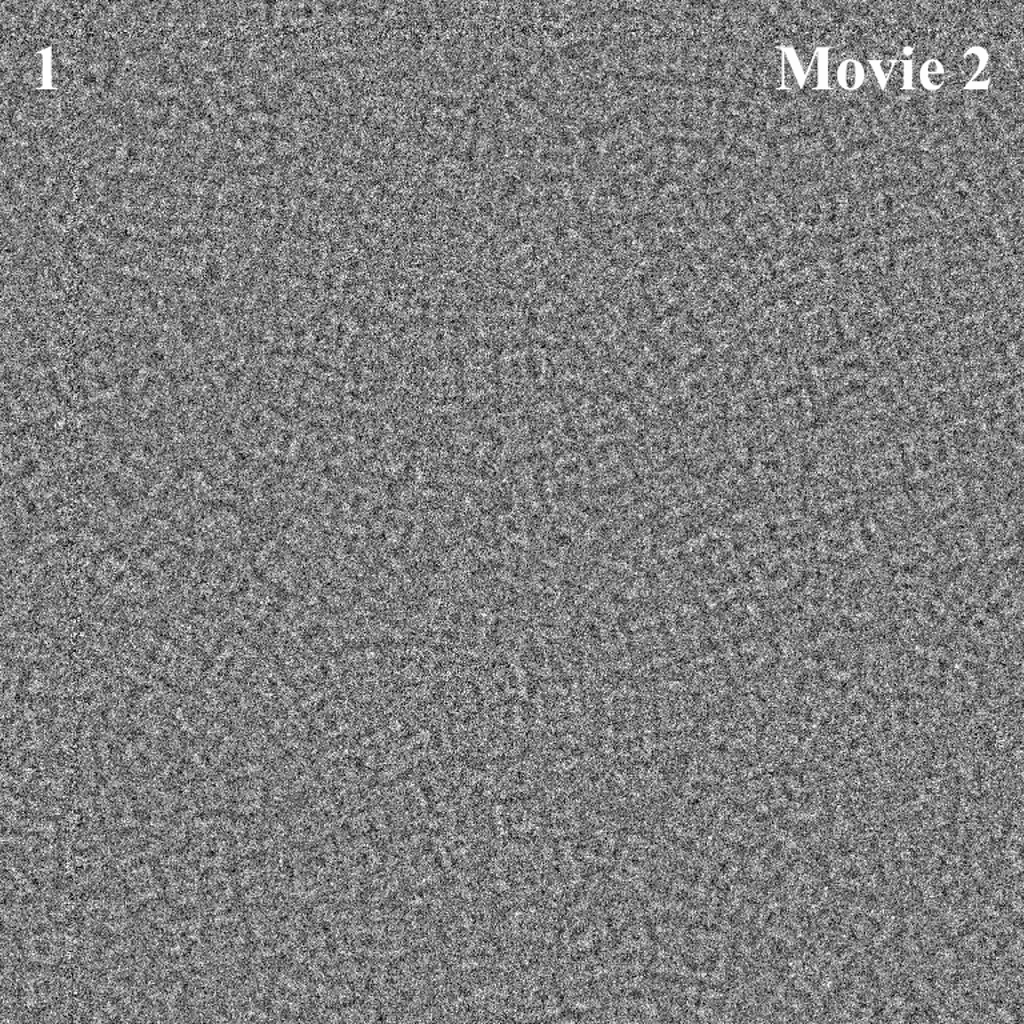

### Movie S3

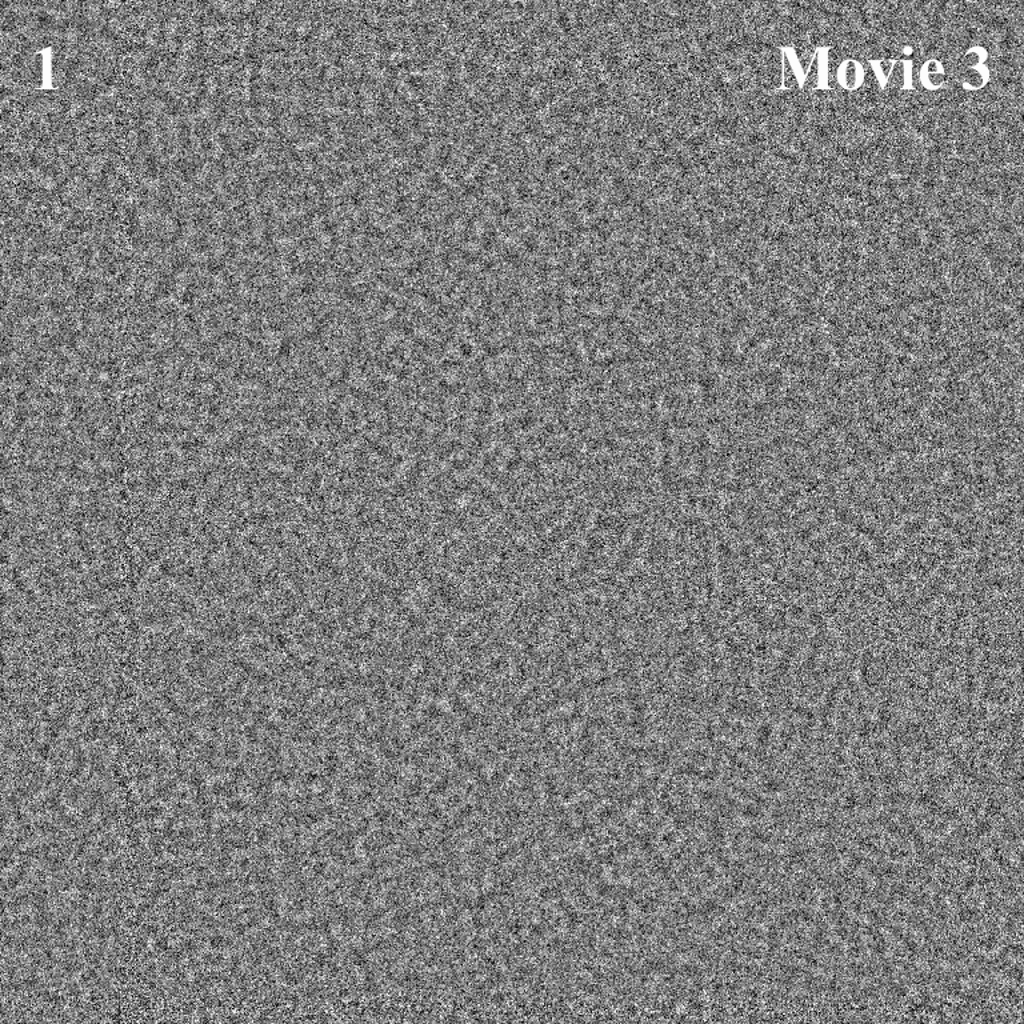
